# Limits to social competence across opposing social roles in a cooperatively breeding cichlid

**DOI:** 10.64898/2026.04.24.719218

**Authors:** A. Ramesh, B. Taborsky

## Abstract

Species in stable social groups engage in diverse social interactions, where social competence - the ability to adjust behaviour using social information - can influence fitness. Yet, whether adaptive behavioural flexibility is expressed across contexts within individuals remains relatively untested. To address this, we exposed cooperatively breeding cichlids (*Neolamprologus pulcher*) to a role-reversal paradigm. In this species, the early social environment shapes social competence, with more competent individuals adjusting behaviour flexibly to social challenges, while individuals also show consistent differences in traits such as aggression. In the present study, individuals were successively assigned to two contrasting roles, smaller territory owners (TOs) and larger intruders (INTs). We predicted role-specific social competence metrics based on behaviours facilitating shelter acquisition. Social competence metrics correlated within, but not across the two roles. Competent TOs showed shorter latencies to submit, more submissive responses to received aggression, and low aggression after initial submission. Competent INTs escalated quickly and relied more on overt aggression rather than displays, allowing faster shelter acquisition. Across roles, individuals competent as TOs were not competent as INTs, and vice versa. This role-specific competence instead appears to be related to aggression across the two roles hinting at the existence of stable behavioural tendencies, which may constrain how social competence shapes behavioural strategies.

## Introduction

Most animals engage in social interactions in diverse contexts over their lifetime. Social competence, that is, the ability of an organism to express appropriate behaviour in response to available social information (Taborsky & Oliveira, 2012), significantly impacts fitness. Initially thought to be a characteristic of complex social systems, the study of social competence has since expanded from humans and primates (Sánchez *et al*., 2001; Kempes *et al*., 2008; van Leeuwen *et al*., 2014; Junge *et al*., 2020) to species exhibiting varying degrees of social complexity (for example, in ravens (Boucherie *et al*., 2019); in fish (Bshary *et al*., 2014; Nyman *et al*., 2018) and in rodents (Williamson *et al*., 2017)). Social competence should benefit most animals as they encounter social challenges during some instances in life, such as competitive interactions, mating or foraging (Taborsky & Oliveira, 2012). Social competence is expected to represent a domain-general social ability and measured via performance traits relating to a clear outcome (Taborsky & Oliveira, 2012). It involves cognitive perception of social situations and decision-making to achieve specific goals while minimizing the costs associated with social interactions (Rose-Krasnor, 1997; Varela *et al*., 2020). For example, if during a territorial contest, an individual perceives a competitor to be unbeatable, either from prior knowledge or from physical differences, and therefore would incur high costs of escalating, it should retreat or submit quickly (McGregor & Peake, 2000; Grosenick *et al*., 2007). But if the opponent is less competitive, then the individual should accurately assess this and escalate competition to secure a resource or reward. This ability of flexibly adjusting behavioural strategies forms the core of social competence.

A conceptual and methodological challenge in the study of social competence is potential circularity. Social competence is often defined post-hoc, only after the behavioural outcome (e.g. achieving a resource) is known. In the previous example, the competent behaviours are classified after observing the outcome of the conflict from the focal individual’s perspective. This risks circularity because the same outcome is then used both to define and to evaluate competence. We therefore require experimental systems in which (1) outcome criteria can be specified a priori, (2) experimental manipulations produce deterministic outcomes, and (3) a priori predictions about competent behaviours can be derived from those known outcomes.

The cooperatively breeding cichlid *Neolamprologus pulcher* is a well-suited model system to meet these requirements. *N. pulcher* forms stable social groups with linear, size-based hierarchies with frequent, well-characterized social interactions, which makes it straightforward to define ecologically relevant social roles and to score role-appropriate behaviours (Jordan *et al*., 2021). Additionally, previous work demonstrates that early-life social environment and predator exposure shape later social performance: individuals reared without parents or conspecifics, with reduced sibling interactions, or under altered predator regimes, show a lowered integration ability into groups and reduced submissive responses when acting as subordinates, thus enabling us to make predictions based on validated foundations (Fischer *et al*., 2015, 2017; Nyman *et al*., 2017; Kasper *et al*., 2019; Camargo-dos-Santos & Taborsky, 2026). Most studies in *N. pulcher* have focused on group-level differences produced by developmental manipulations. Whether competence metrics covary among contexts at the individual level however remains less clear. Therefore, we exposed the same individuals to two opposing social roles within the same social and ecological context of territory intrusion in succession, using a standardized territory intrusion assay (Fischer *et al*., 2015). The assay stages an asymmetric contest between two fish over the ownership of a shelter. Shelters are vital resources in the natural environment of *N. pulcher* as they provide refuge to escape predation and conspecific aggression (Groenewoud *et al*., 2016; Gübel *et al*., 2021). During this test situation, a smaller fish is first established to occupy the shelter and allowed to settle (hereafter “Territory Owner” or TO). Then a much larger “Intruder” (hereafter, INT) is introduced, which strives for the takeover of the territory and shelter from the smaller TO.

This size-asymmetric setup produces highly predictable contest outcomes, with larger intruders typically achieving takeover (as described also in previous studies employing this test, see La Loggia & Taborsky, 2024; La Loggia *et al*., 2024). This predictability allows us to define role-specific, a priori expectations of competent behaviours for each contestant, and to derive operational measures of “social competence” based on efficiency in achieving role-relevant outcomes. Thus behavioural variables were selected a priori to capture role-specific components of contest performance in asymmetric territorial interactions (Table 1). Variable selection was guided by three criteria: (i) ecological relevance to contest outcomes, (ii) expected role-specific contribution to contest efficiency, and (iii) measurability as discrete components of escalation, submission, and aggression regulation. In the territory owner (TO) role, behavioural measures were chosen to quantify the efficiency of transition from territorial defence to submission, as well as post-transition adjustment of aggression in response to the intruder. In the intruder (INT) role, measures quantified the speed and intensity of escalation leading to territory acquisition, and the balance between overt and restrained aggression as alternative escalation strategies (see Table 1).

**Table 1.**
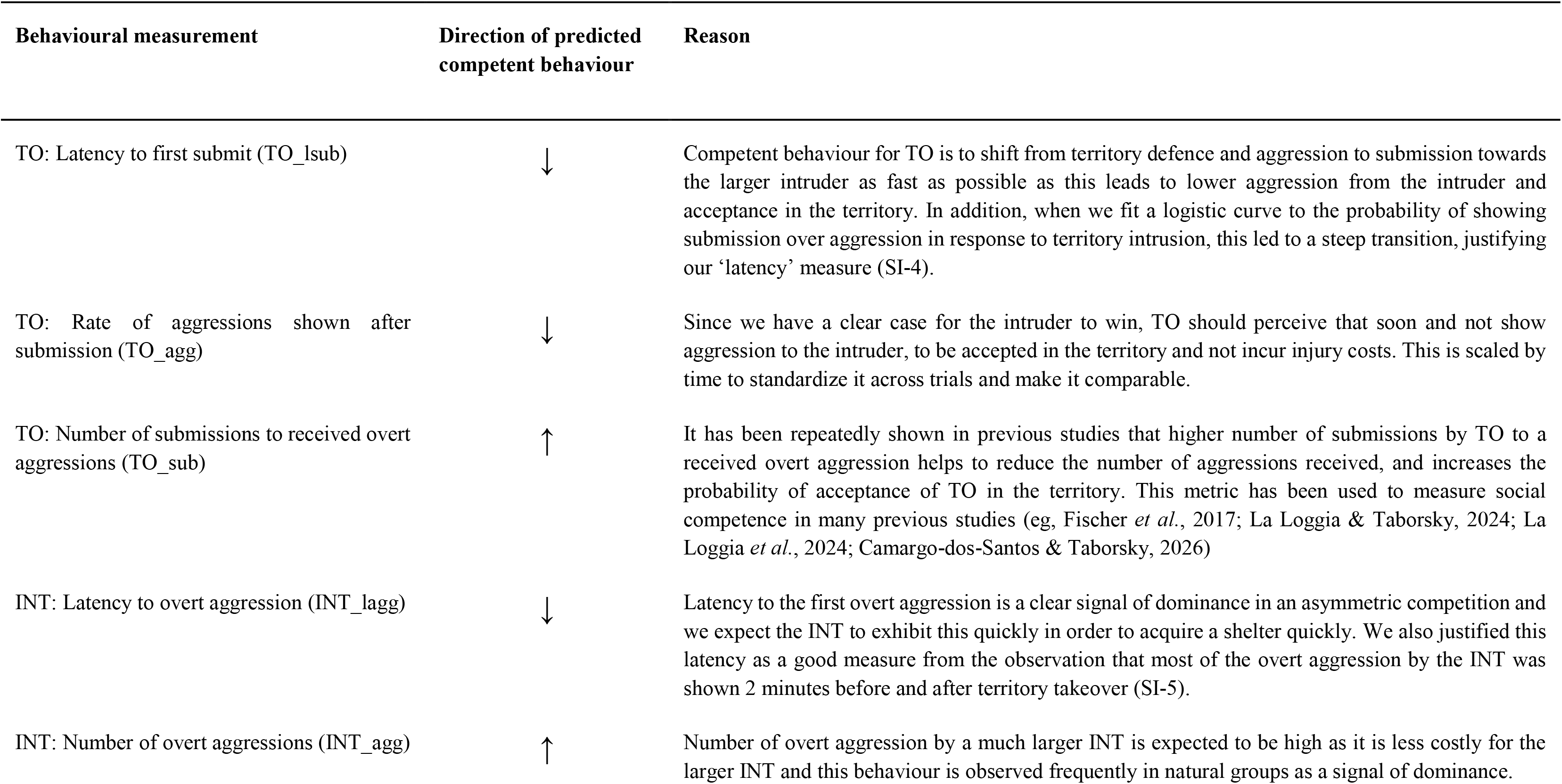
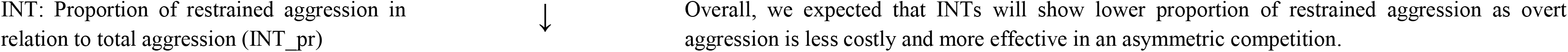
List of the behavioural measurements and their predicted competent levels within the roles of TO and INT.

With this role-reversal design we address 1) whether within a given role, behavioural measures that we label *a priori* as “competent” covary and 2) whether individuals that behave competently in one role also behave competently in the other. We predict that if the behavioural measures used here reflect a common underlying social competence measure, individuals should adjust their behaviour appropriately to each role and show positive correlations in competence within and across contexts. While our present focus is on defining and quantifying behaviours underlying social competence across two roles, linking individual variation in competence to fitness outcomes (survival, reproductive success) remains an important direction for future research.

## Methods

The study was preregistered in OSF (https://doi.org/10.17605/OSF.IO/NPMQ2). Below, we present the description of the methods. We followed the preregistration as closely as possible and noted deviations from preregistration (detailed in SI-1).

### Study species

*Neolamprologus pulcher* is an obligate cooperatively breeding cichlid endemic to Lake Tanganyika, East Africa (Konings, 1998). Social groups in this species typically consist of a dominant breeding pair and 1–25 subordinate individuals, referred to as ‘brood care helpers’ Groups in natural habitats are organized into linear, size-based hierarchies (Dey *et al*., 2013). Helpers engage in vital cooperative tasks such as territory defence, territory maintenance, and alloparental care of eggs and larvae (Taborsky, 2016). Upon maturation, subordinates can either remain within their natal territory as helpers, potentially inheriting the territory in the future, or disperse to obtain a breeding position elsewhere (Stiver *et al*., 2004; Fischer *et al*., 2017; Jungwirth *et al*., 2023). Individuals within a social group interact frequently exhibiting a range of fine-tuned social displays in the context of aggression, submission and affiliation (ethogram described in Reyes-Contreras *et al*., 2019)

### Housing conditions

Experiments were conducted at the Ethologische Station Hasli, University of Bern. Fish were taken from the stock population of the facility, which was bred in the lab over 5-6 generations and was originally derived from Lake Tanganyika from Kasakalawe, Zambia. Housing conditions simulate natural conditions with tap water (total hardness ≥10, carbonate hardness ≥10, pH 7-8) matching Lake Tanganyika water, 13:11 h light:dark cycle with 10 min dimmed light transitions, and water temperature of 27 ± 1 °C. The experimental fish were fed daily after the trials with commercial flakes (5 days/week) and frozen zooplankton (1 day/week). Water changes were done once in 6 weeks, where algae and debris were removed along with 1/4^th^ of the tank water, which was replenished with fresh tap water.

### Experimental procedures

#### Fish selection and housing

Focal fish of standard lengths (SL; from tip of the snout to the end of the caudal peduncle) of 3 – 4 cm (N= 20 males and 20 females) were selected from stock tanks. They were caught in the afternoons on the day before experimentation and housed individually in 50-L tanks provided with a shelter (half of a clay flower pot). The focal fish were randomly assigned to the role of a TO or INT, respectively in the first or in the second test. Stimulus fish of the same sex were selected to be the larger or smaller opponent of the focal fish, depending on the assigned social role of the focal fish. In the case of a focal TO, stimulus INT were chosen to be 0.8 – 1.0 cm larger, and in the case of the focal INT, stimulus TO were 0.8 – 1.0 cm smaller than the focal representing about 25% relative size difference (see La Loggia *et al*., 2024). Six trials were performed daily between May - July 2024. Although winner and loser effects can persist in this system, the effect sizes were demonstrated to be small (Lerena *et al*., 2021) and we intended to minimise this effect by leaving a one-week interval between the role-reversal trials, and the use of novel stimulus fish. Only focal fish were tested in both the TO and INT contexts against a new stimulus fish each time.

#### Territory intrusion assay

When in the role of a TO, focal fish were isolated overnight in the experimental tank. The trials were done in 50-L tanks equipped with a 2-cm layer of sand, a clay flower pot half serving as shelter, and an air stone for oxygen supply. On the day of the trial, we introduced a temporary divider to create a compartment of about 1/4^th^ of the tank space and introduced the INT fish there. After 5 min, the divider was gently removed, releasing the INT. Directly after the release, we recorded all the interactions between the two fish for 20 min using a video camera (Sony camcorder), following the protocol used by La Loggia and colleagues (La Loggia *et al*., 2024). The observer was not present in the room during the recording to avoid disturbance. After the trial, we monitored the status of the fish and aggression for about 5 minutes every hour. At the end of 3 h and again at 4 h after a trial, we scored the TO as being “accepted” or “evicted,” depending on the affiliative and submissive behaviours shown toward the INT, the received and expressed aggression, the distance between the two fish, and whether the TO use the shelter (La Loggia & Taborsky, 2024). A TO was scored as being “accepted” when it showed submissive and affiliative behaviours, stayed close to the larger INT and had access near/to the shelter. In contrast, an “evicted” TO never showed submissive or affiliative behaviours, received aggression from the INT, and was forced to stay in a confined space either near the water surface or in one corner of the experimental tank. At the end of 4 hours, we transferred the fish into their respective individual home tanks.

When in the role of an INT, the focal fish were subjected to the same territory intrusion assay but they now assumed the role of the INT described above, with a smaller stimulus fish adopting the role of an TO.

### Predicted behaviours in the two roles

Video recordings were analysed using BORIS v.6.2.4 (Friard & Gamba, 2016). We scored behaviours over 20 min, classifying them as *restrained aggression* (non-contact displays: head down, frontal approach), *overt aggression* (bites, ramming, chasing, mouth fights), and *submission* (tail quivering) (see SI-2 for more details).

We predicted which correlations among behaviours should occur assuming socially competent behaviour within and across the two roles (Table 1, Fig. 1a). Based on previous studies on contests between size-matched *N. pulcher* (Arnold & Taborsky, 2010), we initially predicted that competent individuals will exhibit more threat displays, as escalation can be costly for both opponents (see preregistration in SI-1). Accordingly, we had hypothesised that successful intruders would show more threat displays and lower levels of overt aggression. We revised this prediction, as the present study involves asymmetric competition with INTs being substantially larger than TOs, and thus the injury cost of quick escalation are minimal for INTs (and very high for TOs). Therefore, in our set-up, overt aggression is expected to be the more efficient and less energetically costly strategy for INTs, as it can reduce the duration of the conflict and enable rapid shelter takeover.

**Figure 1:**
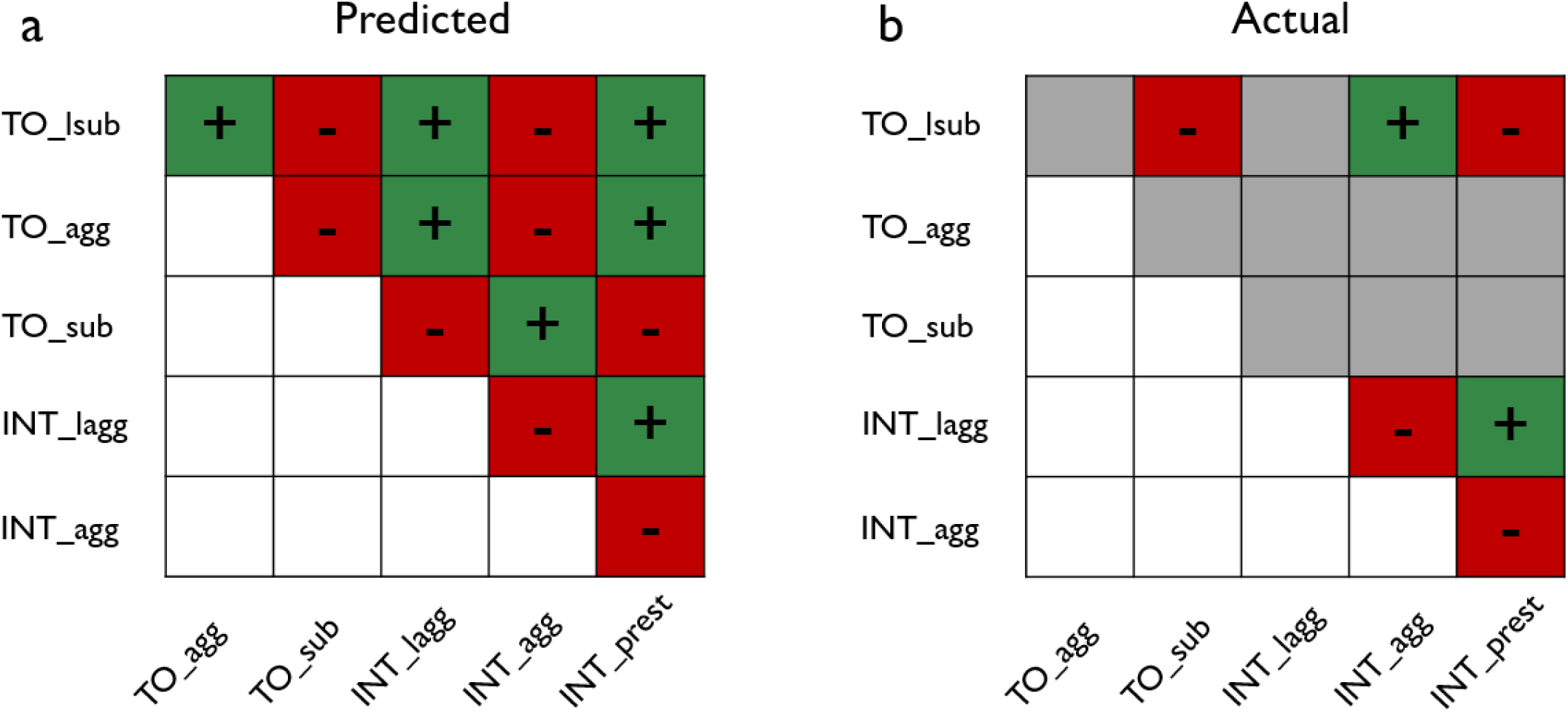
Predicted and actual correlations within and across the two opposing social roles. a. The predicted direction of correlations of different behaviours within roles of TO and INT and across the two roles are derived from Table 1. If the arrows of the two variables that we want to associate point in the same direction in Table 1, a positive correlation is predicted and otherwise a negative one. Positive correlations are depicted by green squares and ‘+’ sign while the negative correlations are represented by red squares and ‘-’ sign. b. Actual correlations obtained from the data. Correlations in colour were significant in at least one of the tests and the grey squares represent non-significant correlations in both the Spearman’s *Rho* and multivariate regression residual correlations (SI-7 & SI-8).

### Predicted outcomes in the two roles

Regarding outcomes, a TO’s success was defined as binary: either being ‘accepted’ in the territory (scored as 1) or ‘evicted’ (scored as 0). For INTs, the successful outcome was a shorter *latency to take over the shelter* (i.e. the latency to occupy the shelter for >5 s). These outcomes for both roles are expected to relate directly to survival in *N.pulcher* in their natural habitat– eviction from a territory substantially reduces chances of survival, and likewise, having no access to a shelter within the safety of a territory increases vulnerability to predation (Heg *et al*., 2004; Groenewoud *et al*., 2016). In addition, behaviours that minimise energetic expenditure and injury costs are expected to be shown.

### Statistical analyses

We first tested how behavioural traits predicted the outcomes *within each context* using regression models, including all fish, both focal and stimulus ones, that were in TO or INT role (N = 81). This was possible because stimulus fish also occupied TO or INT role during the tests, but unlike the focal fish, they were not tested in both the roles. For the TOs, binomial GLMs were fitted with *acceptance state* as response variable; for INTs, we fitted Gaussian GLMs with the *latency to take over the shelter* as response variable. Fixed effects were the behaviour of interest, sex, and stimulus fish aggression. The effect of testing order (TO or INT first) on the latency to submit as TO and latency to overt attack as INT were tested in a linear model and were non-significant, with a low effect size (not included).

To analyse *within- and cross-context* relationships, we used Spearman rank correlations with raw data and calculated residual correlations from Bayesian multivariate models (*brms*) across focal fish tested across both TO and INT roles (N = 42). Latencies were square-root transformed; count data were log-transformed; all variables were scaled to give the z-scores (mean centred and divided by the standard deviation (SD) resulting in mean of 0 and SD of 1). Models included sex and stimulus fish aggression as fixed effects, with weakly informative priors (Lemoine & Lemoine, 2019). Model fit and convergence were checked using standard diagnostics (trace plots, Ȓ, posterior predictive checks). Because each individual was tested once per role, the analyses quantify cross-role associations of social competence measures but do not estimate repeatability or behavioural consistency in a quantitative personality framework. All analyses were done in R (v4.3.2) with *lme4* (Bates *et al*., 2015a), *brms* (Bürkner, 2017), *DHARMa* (Hartig, 2024), and *ggplot2* (Wickham, 2016) packages.

Detailed description of the statistical analyses is given in SI-3.

## Results

### Temporal structure of contest dynamics

Direct interactions were essential for the switch in behaviour from aggression to submission for territory owner (TO) and territory takeover for intruder (INT). Direct interactions between contestants were concentrated around the transition points of the contest, which was the initial territory takeover / territory loss. For TOs, submission was typically preceded by overt aggression from the intruder, with the probability of submission increasing sharply following overt aggressive interactions with INT (SI-4). For intruders (INTs), most overt aggression occurred within the two minutes preceding and following shelter acquisition (SI-5), indicating that escalation was closely associated with the transition in territory ownership.

Nevertheless, aggressive and submissive interactions frequently continued after takeover, suggesting that social interactions persisted beyond the initial resolution of territory ownership.

### Social competence within roles

When in the role of a TO, fish were evicted in 25 out of 81 trials. Fish, which submitted sooner, had a higher probability of being accepted (Fig 2a, Table 2). The probability of being accepted was not predicted by variation in aggression after submission and submissions per received aggression (Fig 2b, c, Table 2). When in the role of an INT, faster shelter takeover was linked to showing overt attacks sooner, expressing more overt aggression, and using relatively fewer restrained displays (Fig. 1, Table 2).

**Figure 2:**
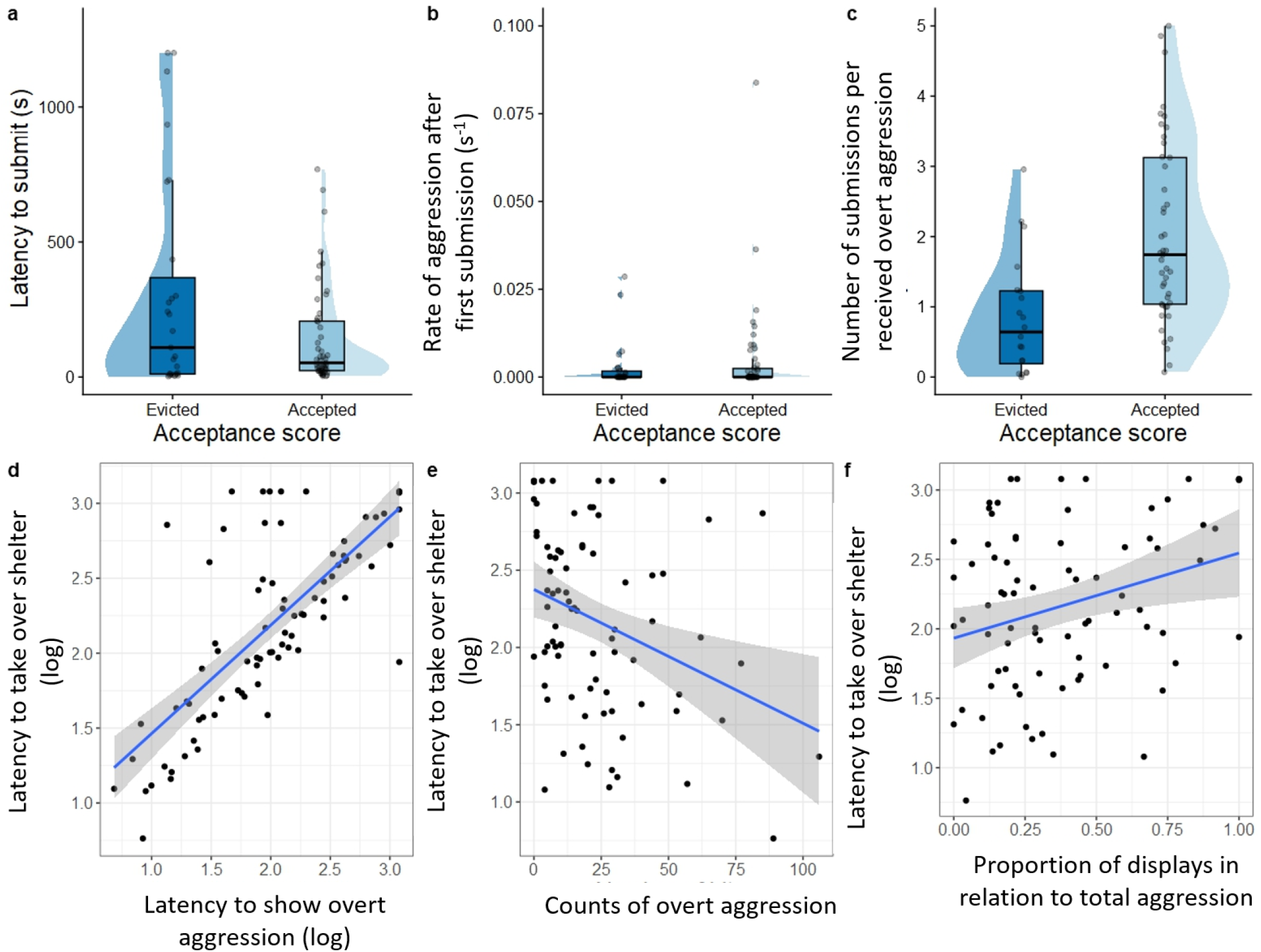
Relationship of behaviours with the outcome in each role. a, b and c show the differences in behaviours between accepted and evicted fish in TO. d, e and f show the relationship between the behaviours and the latency to take over the shelter (log10-transformed) in INT. (N = 81).

**Table 2.** Summary of linear models of behavioural traits related to the outcome within each context. The different behaviours of interest, ‘stimulus fish aggression’ and ‘sex’ were fitted as fixed effects. Estimates of fixed effects (β) are given with their 95% confidence intervals (CI) and the variance components of random effects. Significant effects are denoted in bold (N = 81).

| *Response variable* | **Role - TO** |  |  | **Role - INT** |  |  |
| --- | --- | --- | --- | --- | --- | --- |
|  | *Probability to be accepted in the shelter* |  |  | *Latency to take over the shelter* |  |  |
| **Behaviour of interest** | **Latency to submit** | **Rate of aggressions after submission** | **No. submissions to received aggression** | **Latency to bite** | **Total number of bites** | **Proportion of restrained aggression** |
| *Fixed effects* | $\beta$ (95% CI) | $\beta$ (95% CI) | $\beta$ (95% CI) | $\beta$ (95% CI) | $\beta$ (95% CI) | $\beta$ (95% CI) |
| **Intercept** | **1.41**<br>(0.30, 2.65) | **0.81**<br>(-0.12, 1.79) | 1.12<br>(0.03, 2.29) | 53.08<br>(-41.21, 14.38) | **291.88**<br>(161.39, 422.37) | -13.38<br>(-152.10, 125.34) |
| **Effect of behaviour of interest** | **-0.002**<br>(-0.004, 0.00) | 15.60<br>(-28.92, 88.94) | -0.02<br>(-0.11, 0.09) | **0.63**<br>(0.45, 0.81) | **-4.06**<br>(-7.02, -1.10) | **449.46**<br>(210.94, 687.97) |
| **Stimulus fish aggression** | -0.01<br>(-0.03, 0.01) | 0.00<br>(-0.03, 0.01) | -0.01<br>(-0.04, 0.01) | **8.84**<br>(5.76, 11.91) | **11.49**<br>(7.79, 15.20) | **11.18**<br>(7.72, 14.64) |
| **Sex M** | -0.27<br>(-1.36, 0.77) | -0.04<br>(-1.04, 0.94) | -1.11<br>(-1.14, 0.90) | 89.13<br>(-21.40, 199.67) | 41.26<br>(-94.86, 177.38) | 115.72<br>(-13.18, 244.61) |
| **Residual variance** | 4.85 | 4.59 | 4.43 | 5.61 x 10<sup>4</sup> | 8.44 x 10<sup>4</sup> | 7.40 x 10<sup>4</sup> |

Within each social role, several measures covaried in the predicted directions (Fig. 1a & b), and these associations were supported by both raw correlations and multivariate residual correlations (SI-7 & SI-8).

### Social competence across roles

Across roles, the outcome measures, that is the probability of being accepted as a TO and latency to shelter takeover as an INT were uncorrelated (SI-6). Thus, individuals that were more likely to be accepted in the TO role did not show faster shelter takeover in the INT role.

Behavioural measures also showed weak and inconsistent associations across the two opposing roles (Fig. 1b). Raw correlations indicated moderate relationship between TO latency to submit and INT aggression measures (overt aggression: ρ = 0.35, p = 0.03; proportion of restrained aggression: ρ = –0.36, p = 0.03; Suppl. Info. 7). However, these associations were not supported by multivariate residual correlations (Suppl. Info. 8).

## Discussion

This is the first study to operationalise and test behavioural measures intended to reflect social competence across two opposing roles in a role-reversal task. We expected these measures to covary both within and across roles if they captured a common underlying component of social competence as a general social ability. Specifically, we expected the measures of social competence in the smaller territory owner (TO) and larger intruder (INT) roles during an asymmetric competition to covary. Indeed, some of the behavioural measures of social competence were correlated, and related to the appropriate outcome, when considered within a given role. However, individuals that had positive outcomes in one role did not necessarily have this also in the other role (SI-6). Competence metrics were not related across social roles (SI-7 & SI-8) and instead, we observed a positive association in overt aggression across roles (SI-9), suggesting some consistency in aggressive behavioural tendencies across the two roles. Overall, cross-role associations among the behavioural measures were weak or absent. Importantly, these results do not provide evidence for a general social competence trait expressed consistently across roles, and neither do they exclude the possibility that some stable behavioural tendencies influence performance across social roles.

### Limits to behavioural flexibility

Individual tendencies, also referred to as animal personalities, may constrain behavioural flexibility and hence social competence (Sih *et al*., 2004; Dall *et al*., 2012). Previous studies demonstrated that *N. pulcher* individuals exhibit consistent differences in a range of behavioural traits, and showed that aggression and submission are repeatable (La Loggia et al. 2024) and that aggression, boldness and explorative behaviour form a repeatable syndrome in this species (Chervet *et al*., 2011). Such stable tendencies may influence how individuals express behaviour across different social contexts, potentially limiting behavioural flexibility at the short timescale relevant for contests. In *N. pulcher*, consistent individual differences in social behaviour have also been linked to helping propensity in subordinate individuals (Riebli *et al*., 2012, Fischer et al. 2017). Likewise, high aggressive, risk-prone or active individuals showed higher helping tendencies (Le Vin *et al*., 2011; Kasper *et al*., 2019). These patterns suggest that social behaviour may be influenced by partially shared physiological or neuroendocrine mechanisms, including stress-axis regulation (Reddon *et al*., 2014; Stettler *et al*., 2021; Antunes *et al*., 2022, 2024). Taken together, our results are consistent with an interaction between role-specific performance and consistent individual differences in behaviour.

### Group-level vs individual-level differences

Previous work in *N. pulcher* have repeatedly shown that early social environment produces consistent between-group differences in social competence and related performance measures such as conflict resolution duration (Fischer *et al*., 2015, 2017; Kasper *et al*., 2019; La Loggia & Taborsky, 2024; La Loggia *et al*., 2024). In contrast, our context-specific metrics do not show a corresponding across-role correlation at the individual level.

This difference likely reflects both the experimental design and level of analysis. While previous studies focussed on differences between different developmental treatment groups, our study focused on individuals raised under broadly comparable laboratory conditions, thereby focusing on within-group individual variation. As a result, group-level differences observed in earlier work do not necessarily imply corresponding associations at the individual-level.

Such divergences between levels of analysis are well-known in the behavioural and social sciences: correlations estimated at an aggregate or group level can differ or even have opposite signs from individual-level correlations (Robinson, 1950; Subramanian *et al*., 2009), leading to a pattern depicted in SI-9. Group-level data can mask or smooth individual heterogeneity (e.g., unmeasured traits, behavioural syndromes), producing a group mean relationship that does not hold at the individual-level. Further, processes that create group-level differences, such as developmental canalization, are not the same mechanisms that generate within-individual behavioural variation. Individual behaviours may instead be shaped by state-dependent feedbacks, short-term physiological shifts, or context-dependent plasticity and trade-offs (specialization).

### Limitations of the study

A key limitation of the present study is that each individual was tested once per social role. As a consequence, our analyses quantify cross-role associations among behavioural measures, but they do not allow estimation of behavioural repeatability or the partitioning of within-and among-individual variance that is required in a quantitative personality framework (Niemelä & Dingemanse, 2018; Dingemanse & Wright, 2020; Barbosa & Morrissey, 2021). The current study was not designed to assess behavioural consistency or personality structure, but to test whether behavioural measures intended to reflect competent performance are expressed in a correlated manner across opposing social roles within the same individuals. Within this framework, the absence of strong cross-role associations may reflect substantial within-individual variability, role-dependent behavioural expression, and potentially stable individual differences that are expressed across roles. Clarifying the extent to which these mechanisms contribute will require repeated measurements of the same individuals across roles and contexts.

## Conclusions

Measuring social competence is challenging because it is a multidimensional trait expressed through flexible behaviours that vary with social context. Our study tested whether behavioural measures intended to capture social competence covary across opposing social roles within the same individuals. We found limited cross-role covariation, suggesting that role-specific behavioural expression plays an important part in shaping performance in asymmetric social interactions.

More broadly, our results indicate that group-level differences arising from early-life experience do not necessarily translate into corresponding patterns of individual-level covariation across roles and contexts. We therefore emphasize the importance of assessing behavioural performance across multiple roles and contexts within individuals while distinguishing between cross-context covariation and behavioural repeatability as distinct analytical targets. Doing so will help clarifying how social competence and consistent behavioural types interact in maintaining variation in animal social behaviour.

## Ethical note

All experiments were conducted at the Ethological Station Hasli of the Institute of Ecology and Evolution of the University of Bern, Switzerland. The captive holding of the fish was carried out under licence no. BE 4/22 and experiments were conducted under licence no. BE 34/22 of the Veterinary Office of the Canton of Bern. In addition, we followed the guidelines provided by the Association for the Study of Animal Behaviour for the use of animals in behavioural research (ASAB Ethical Committee & ABS Animal Care Committee 2022). During housing, all fish were monitored daily during feeding by noting normal feeding behaviour, posture, morphology and behaviour. Evicted fish, recognised by limited movement and increased aggression by conspecifics were identified during daily monitoring and immediately isolated. During the contests, the size difference between the fish (about 25%) ensured that escalated aggression and mouth-fights did not occur, which might occur in more similarly sized fish with similar competitive abilities. After the 20 minute recording, fish were observed for 5-10 minutes every hour for 4 hours. These were similar to husbandry and group establishment protocols established in this system. All fish were retained in our permanent breeding stock after the end of the study.

## Data availability statement

The raw and secondary datasets and all R codes used in this paper are available as supplementary data. They will be uploaded to a Zenodo repository if accepted.

## Funding

This work is supported by the Swiss National Science Foundation (SNSF) grant (grant number: 207448) to BT, through which AR is employed.

## Supporting information

R code for plots and analysis - revised

## Supplementary information for the paper titled “Limits to social competence across opposing social roles in a cooperatively breeding cichlid”

### Supplementary information 1: Changes to preregistration

We followed the preregistration as closely as possible. Below, we explicitly distinguish between our preregistration and subsequent revisions.

#### 1. Revision of predictions regarding INT overt aggression behaviour

The preregistration assumed symmetric contests between size-matched individuals and predicted that successful intruders would exhibit higher levels of threat displays and lower levels of overt aggression, as escalation was expected to be costly for both opponents. After conducting a pilot study, we identified a theoretical mismatch between this assumption and our experimental design. Specifically, our study involves asymmetric contests in which intruders (INTs) are substantially larger than territory owners (TOs), to ensure that INTs always win the contest for shelter. Under such conditions, the costs of escalation are reduced for INTs and substantially increased for TOs. Based on contest theory for asymmetric interactions (Hammerstein, 1981; Leimar & Enquist, 1984), we therefore revised our prediction: successful intruders are expected to exhibit higher levels of overt aggression, as this strategy is likely to reduce contest duration and facilitate rapid territory acquisition.

#### 2. Omission of repeated trials

The preregistration planned two repeated trials per individual in each social role to assess behavioural consistency. We omitted this component because the primary aim of the study was to test whether behavioural measures of social performance covary across opposing social roles within individuals, rather than to estimate repeatability or among-individual variance in a personality framework. Addressing repeatability would require substantially more repeated measurements per individual than feasible within the constraints of the experimental design, and limited repetition would not have provided reliable estimates of consistency (Wolak *et al*., 2012). This modification reflects a refinement of the study’s inferential scope rather than a change to the primary hypotheses.

**Supplementary information 2:**
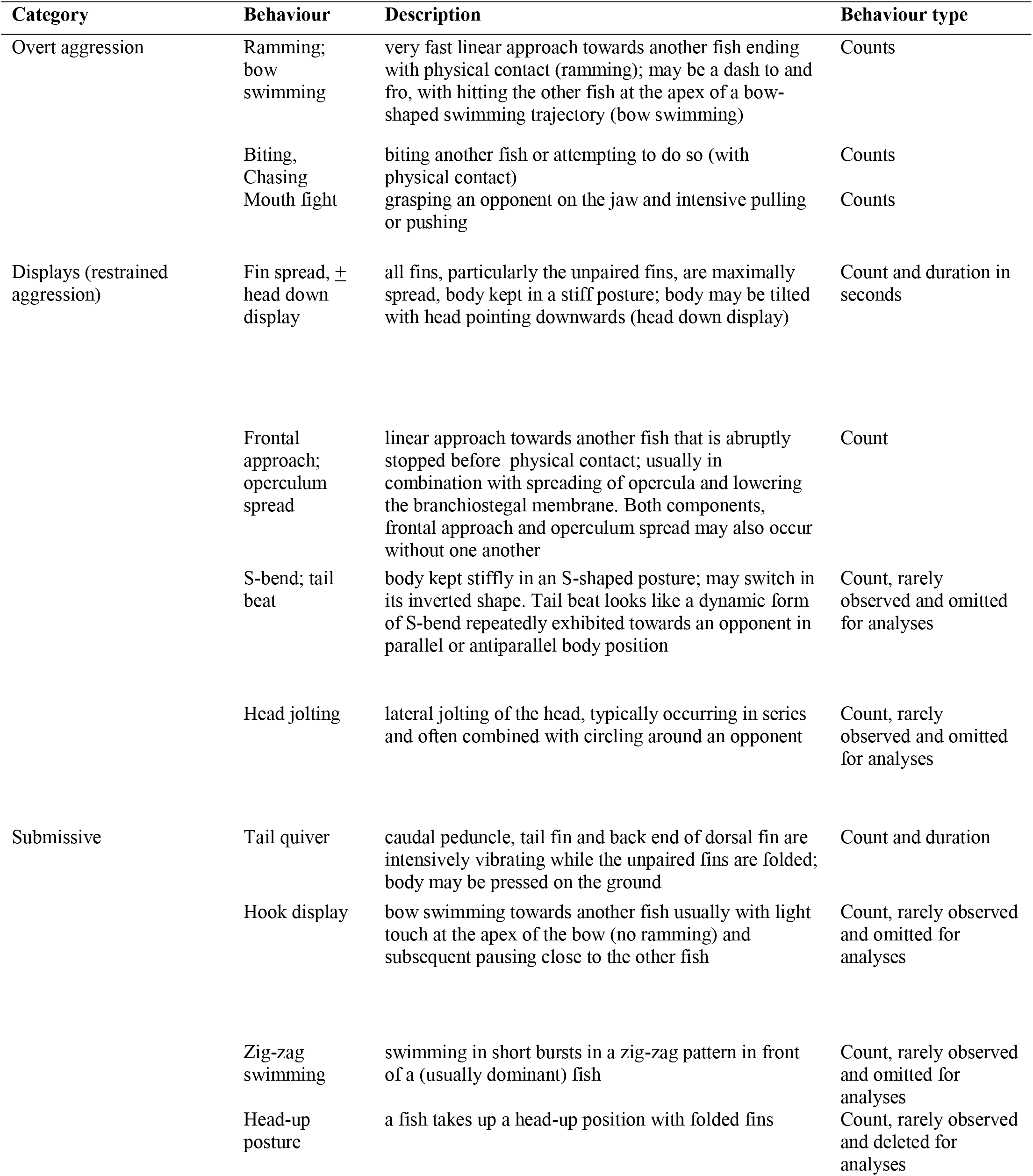
List of behaviours scored and their description.

| Category | Behaviour | Description | Behaviour type |
| --- | --- | --- | --- |
| Overt aggression | Ramming; bow swimming | very fast linear approach towards another fish ending with physical contact (ramming); may be a dash to and fro, with hitting the other fish at the apex of a bow-shaped swimming trajectory (bow swimming) | Counts |
|  | Biting, Chasing | biting another fish or attempting to do so (with physical contact) | Counts |
|  | Mouth fight | grasping an opponent on the jaw and intensive pulling or pushing | Counts |
| Displays (restrained aggression) | Fin spread, ± head down display | all fins, particularly the unpaired fins, are maximally spread, body kept in a stiff posture; body may be tilted with head pointing downwards (head down display) | Count and duration in seconds |
|  | Frontal approach; operculum spread | linear approach towards another fish that is abruptly stopped before physical contact; usually in combination with spreading of opercula and lowering the branchiostegal membrane. Both components, frontal approach and operculum spread may also occur without one another | Count |
|  | S-bend; tail beat | body kept stiffly in an S-shaped posture; may switch in its inverted shape. Tail beat looks like a dynamic form of S-bend repeatedly exhibited towards an opponent in parallel or antiparallel body position | Count, rarely observed and omitted for analyses |
|  | Head jolting | lateral jolting of the head, typically occurring in series and often combined with circling around an opponent | Count, rarely observed and omitted for analyses |
| Submissive | Tail quiver | caudal peduncle, tail fin and back end of dorsal fin are intensively vibrating while the unpaired fins are folded; body may be pressed on the ground | Count and duration |
|  | Hook display | bow swimming towards another fish usually with light touch at the apex of the bow (no ramming) and subsequent pausing close to the other fish | Count, rarely observed and omitted for analyses |
|  | Zig-zag swimming | swimming in short bursts in a zig-zag pattern in front of a (usually dominant) fish | Count, rarely observed and omitted for analyses |
|  | Head-up posture | a fish takes up a head-up position with folded fins | Count, rarely observed and deleted for analyses |

### Supplementary information 3: Detailed description of the statistical methods and justification for using both correlation and multivariate modelling approaches for between-context correlation estimates.

#### 1. Relationship between behavioural traits and outcome variable within each context

To test the relationship between our behavioural traits of interest and positive outcomes within each context, we first computed linear regression models for each of the behavioural traits in separate regression models with the positive outcome as the response variable (‘Binary acceptance score’ for TO and ‘Latency to take over the shelter’ for INT contexts) and ‘Behaviour of interest’, ‘Sex’, ‘Stimulus fish behaviour (overt aggression)’ as the fixed effects. The size differences between focal and stimulus fish can affect their interaction and therefore the outcome. However, the differences between SLwere maintained between 8 – 10 mm (with 4 cases having 7, 11, 12 and 14 mm difference in SL) and therefore size was not included in any of the models. Generalised linear models were fitted with Binomial error distribution with ‘logit’ link for ‘Binary acceptance score’ in the context of TO and Gaussian errors for ‘Latency to take over the shelter’ as INT.

In order to further understand whether the behaviours were different for different sexes, we fitted another set of generalized linear models with ‘Behaviour of interest’ as the response variable and ‘Sex’ and ‘Stimulus fish behaviour (overt aggression)’ as the fixed effects. The models were fitted with Gaussian errors for ‘Latency to submit’, ‘Latency to bite’ and ‘Ratio of displays to total aggression’. The ‘Number of submissions’, ‘Number of bites after first submission per time’ and ‘Number of bites to TO’ are fitted with a generalized linear mixed model with Poisson error and OLRE to account for over dispersion.

All models were fitted using the ‘*glm*’ function of the ‘lme4’ package (Bates *et al*., 2015b). The statistical significance of fixed effects was assessed based on the 95% confidence interval (CI): an effect was considered significant when its 95% CI did not include zero. Model fits were checked by plotting residuals in a Q-Q plot and visually analysing their fit to normality. For Binomial and Poisson models, overdispersion was checked by using the ‘DHARMa’ package (Hartig, 2024).

#### 2. Relationship of behavioural traits within and across contexts

To analyse the relationship among behavioural traits within and across the two contexts, we used two complementary methods to cross-check our analyses.

##### 1. Correlations

We analysed the pair-wise relationship among the different behaviours by Spearman rank correlations. Since the focal fish were tested in both contexts, this was the most straightforward way to analyse the relationship across contexts. Rank correlations give a good indication of the direction of the effects without the need for transforming the data to fit a particular distribution. However, they cannot account for other effects affecting the data such as ‘Sex’ and ‘Stimulus fish behaviour (overt aggression)’ and they also cannot estimate the variance and covariance of all the behaviours together. We computed pair-wise correlations within and across contexts using Spearman correlations in ‘base R’.

##### 2. Multivariate models

Multivariate models are a robust method for analyzing the variance-covariance among multiple variables simultaneously. However, we are limited by the fact that only multivariate Gaussian distributions can be fitted. We therefore square-root transformed the latencies and log-transformed the count data and scaled and centered all variables to zero, in order to fit them into multivariate regression with multivariate Gaussian errors. We fitted and ‘Sex’ and ‘Stimulus fish behaviour (overt aggression)’ as the fixed effects in Bayesian multivariate response models using the ‘brms’ package (v2.18.0; (Bürkner, 2017)). For the ‘fixed’ effects, we used ‘weakly informative’ Gaussian priors (mean = 0, SD = 1) (Lemoine, 2019) . For the estimating the residual correlation matrix, we used the default priors of brms (LKJ(1)). We ran four chains with 1000 discarded warmup iterations, followed by 3000 sampling iterations, thus yielding 12000 posterior samples per model. Trace plots were used to monitor proper mixing of chains. Convergence of chains was ensured by verifying that *Ȓ* values were close to 1.00 (not greater than 1.01). Model fits were evaluated by inspecting posterior predictive checks, using the pp_check() and bayes_R2() function of brms.

Ultimately, we evaluated whether the two methodologies aligned in terms of their effects and effect sizes for interpreting our results. Since we used disjunction testing (which necessitated both analyses to yield the same result and interpretation for a definitive effect) and individual testing (to examine the relationship between each of our behavioural measures and the positive outcome), we opted to not use any correction for multiple testing (Rubin, 2021).

**Supplementary information 4:**
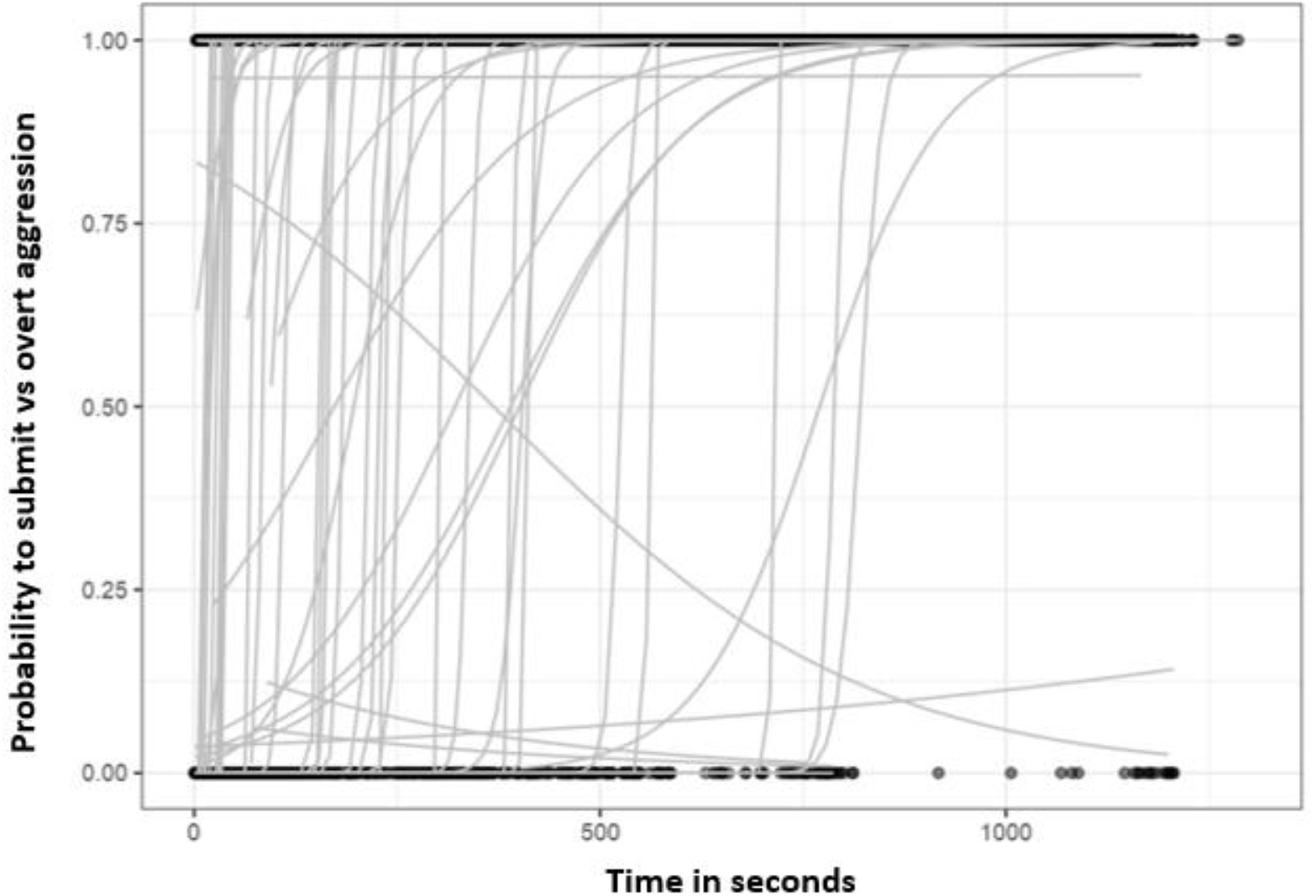
Dynamics of submission behaviour by TO. This figure shows changes in the probability of submission relative to overt aggression over time for territory owners (TOs). Territory owners first respond to a territory invasion by aggressive responses before transitioning to submission when intruders are larger. Binomial logistic curves were fitted for each individual, with submission coded as 1 and overt aggression as 0. Most individuals showed an increase in the predicted probability of submission over time, indicating a transition from predominantly aggressive to submissive behaviour, whereas five individuals showed the opposite pattern. The steep slopes observed for most curves indicate that this behavioural transition occurred over a relatively short time period.

**Supplementary information 5:**
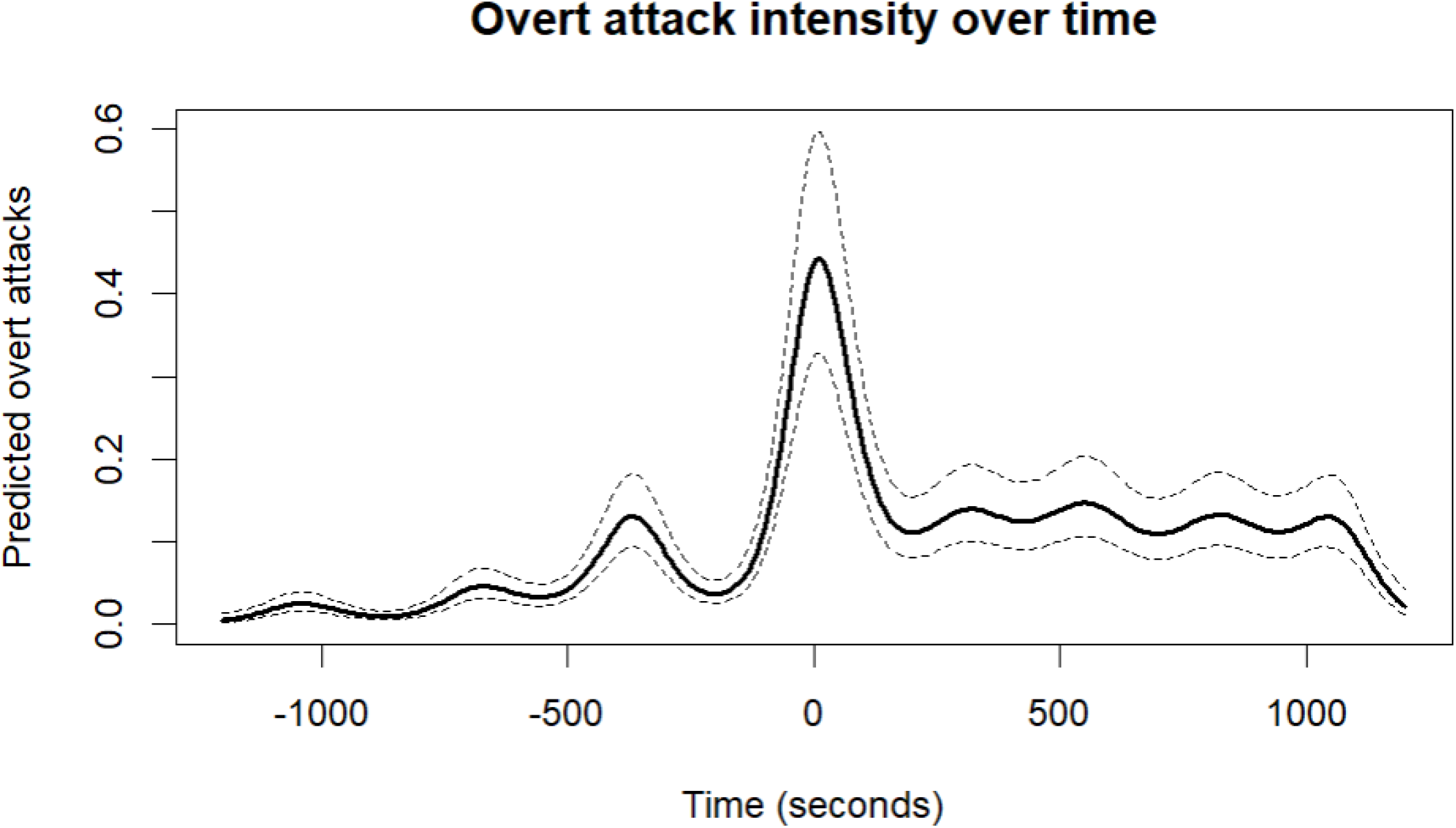
Dynamics of overt aggression behaviour by INT. This figure shows the temporal dynamics of aggressive behaviour by intruders (INT) relative to shelter takeover, with time centered at zero for each individual at the moment of takeover. The x-axis thus runs from -20 minutes to +20 minutes. Aggression peaks during takeover and within approximately two minutes post-takeover, followed by a lower level of aggression over the remainder of the trial. This pattern is consistent with an initial phase of contest escalation and immediate post-takeover dominance consolidation, followed by reduced aggression as interactions stabilize. Temporal dynamics were quantified using a generalized additive model (GAM) fitted with a negative binomial error distribution and log link to account for overdispersed count data (n = 19,039 observations). The model included a smooth term for time since takeover in 10s windows (*s(time10), k = 20; edf = 17.46, Ref.df = 18.65, χ² = 620.6, p < 2 × 10⁻¹⁶*) and a random intercept for individual identity (*s(Observation.id, bs = “re”); edf = 71.37, Ref.df = 78.00, χ² = 895.1, p < 2 × 10⁻¹⁶*). Predicted values represent the population-level smooth over time, with random effects held constant, highlighting a clear transient peak in aggression around the time of takeover.

**Supplementary information 6:**
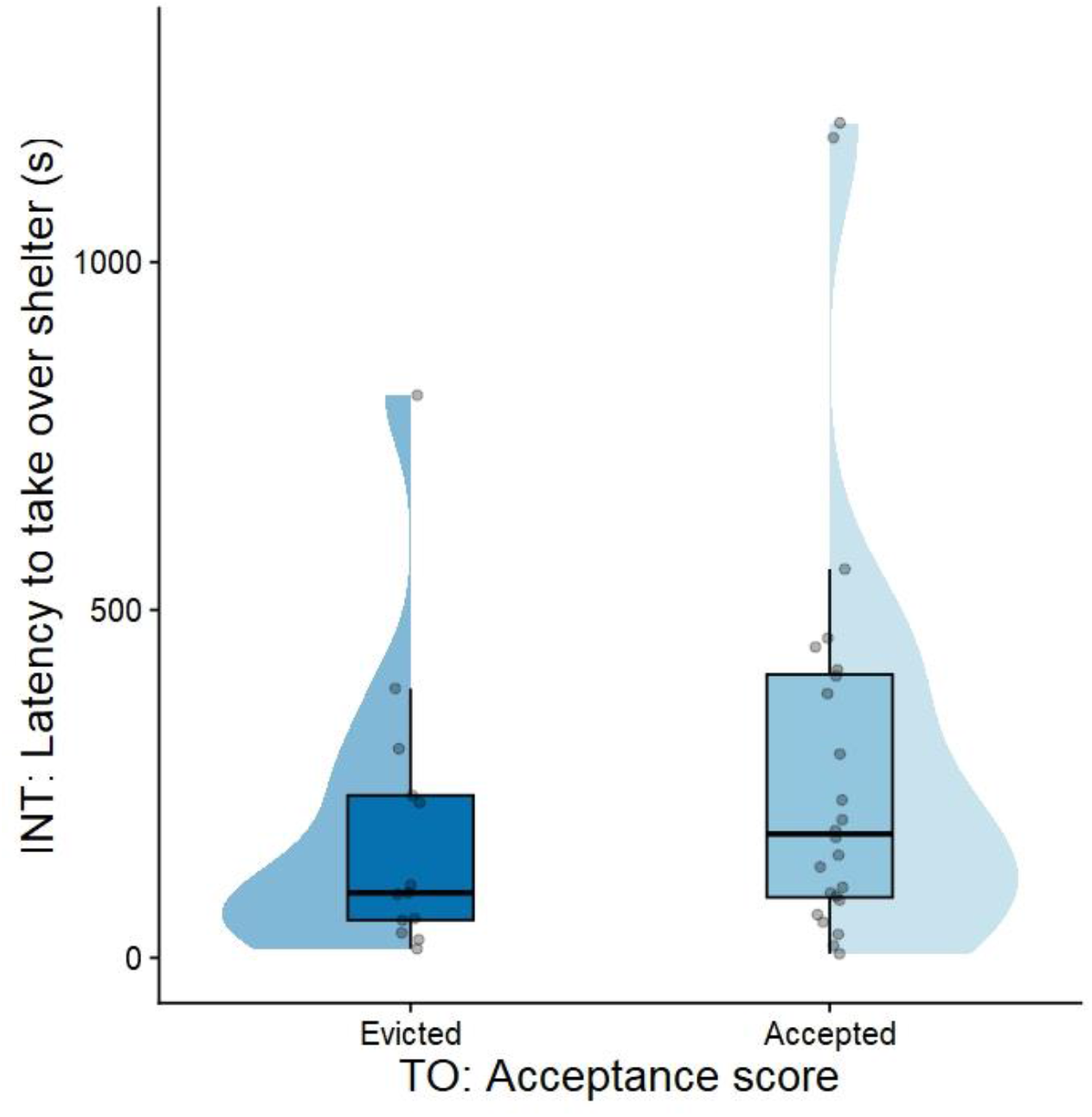
Relationship between the two outcome variables in the TO and INT context. Here, we show the difference between the accepted and evicted fish (positive outcome as TO) and the latency to take over the shelter (positive outcome as INT) for the same focal fish. The differences were not statistically significant. (Wilcox test: W = 190.5, p-value = 0.2794)

**Supplementary information 7:**
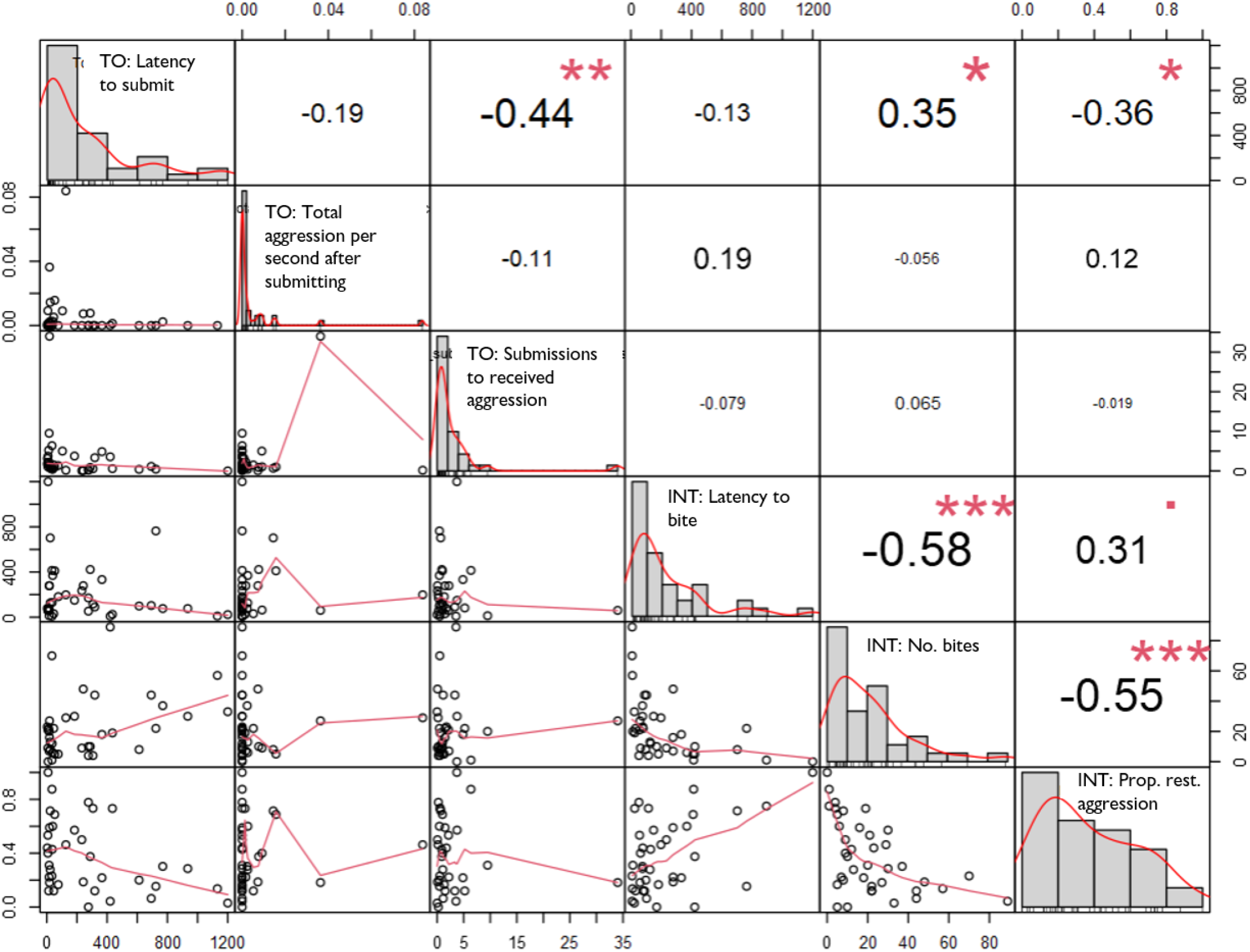
Spearman correlation matrix for pairwise raw correlations within and across contexts. We see that within the context of TO, only Latency to submit and Submissions to received aggression are correlated while in the context of INT, all three behaviours are correlated. We also find that the TO: Latency to submit correlated positively to INT: Number of overt aggression, and negatively to the proportion of restrained aggression (which is related to the overt aggression by setting).

**Supplementary information 8:**
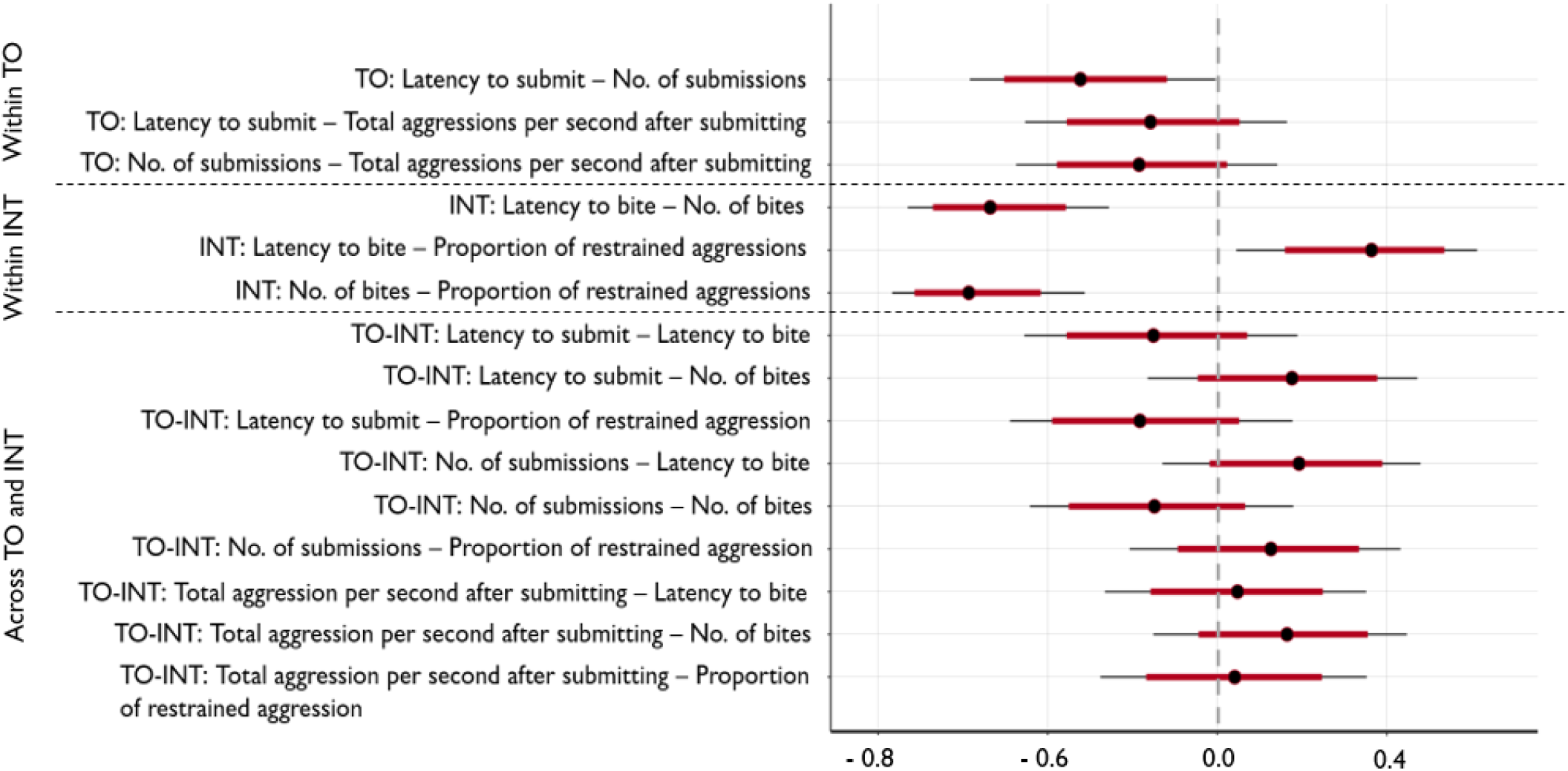
Within and across context residual correlation estimates with 95% HPD (highest probability density) from multivariate analyses. The figure shows the residual correlation model outcome of multivariate regression among the different behavioural traits of interest. Credible intervals level: 0.8 (80% intervals, red thick line) outer_level: 0.95 (95% intervals). When the intervals do not overlap with 0, they can be interpreted as a ‘significant’ effect similar to frequentist statistics. Within the TO role, latency to submit was correlated with the number of submissions, after controlling for the aggression of the interacting partner. Within INT, the latency to first overt aggression was correlated with the number of overt aggressions and the proportion of restrained aggression. The other correlations were not significant.

**Supplementary information 9:**
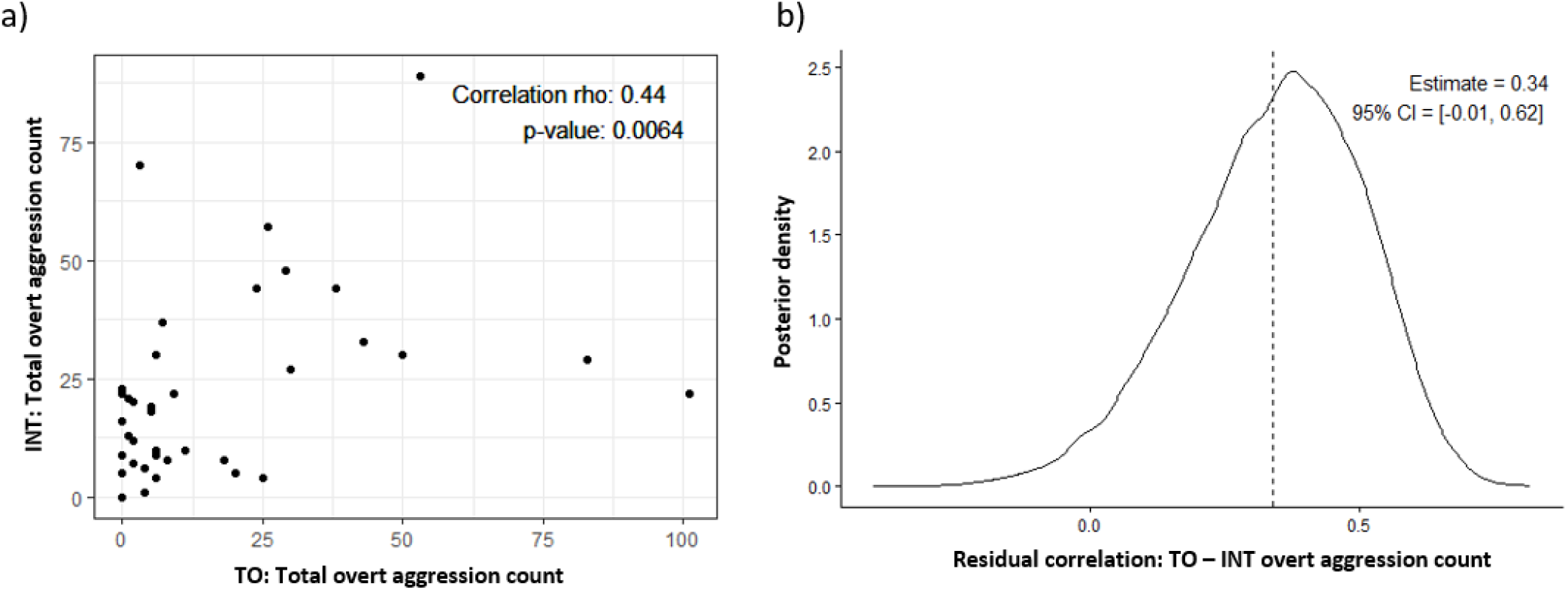
Overt aggression expressed by the focal individual in the role of TO positively correlated with overt aggression expressed in the role of INT. a) Spearman rank correlation of total overt aggression (bites) expressed by the same focal individual across the two roles show a positive correlation indicating that individuals that are more likely to be aggressive in one role are also more aggressive in the other role. However, this could also be masked due to the effect of the interacting partner. Therefore, the same analysis was run as a multivariate analysis, controlling for the overt aggression expressed by the interacting partner in the two roles. b) Bayesian estimate of across role correlation of overt aggression in the form of bites. Posterior distribution of the residual correlation between individual INT and TO bite counts estimated using a multivariate Gaussian model in `brms`. Counts were log-transformed and scaled prior to analysis. The model included partner behaviour as predictors for each role and estimated the residual correlation between the responses of the focal individual in the INT and TO roles. The dashed line indicates the posterior mean residual correlation of ρ = 0.34 (95% Credible Interval = −0.01, 0.62), after taking the partner behaviour into account.

**Supplementary information 10:**
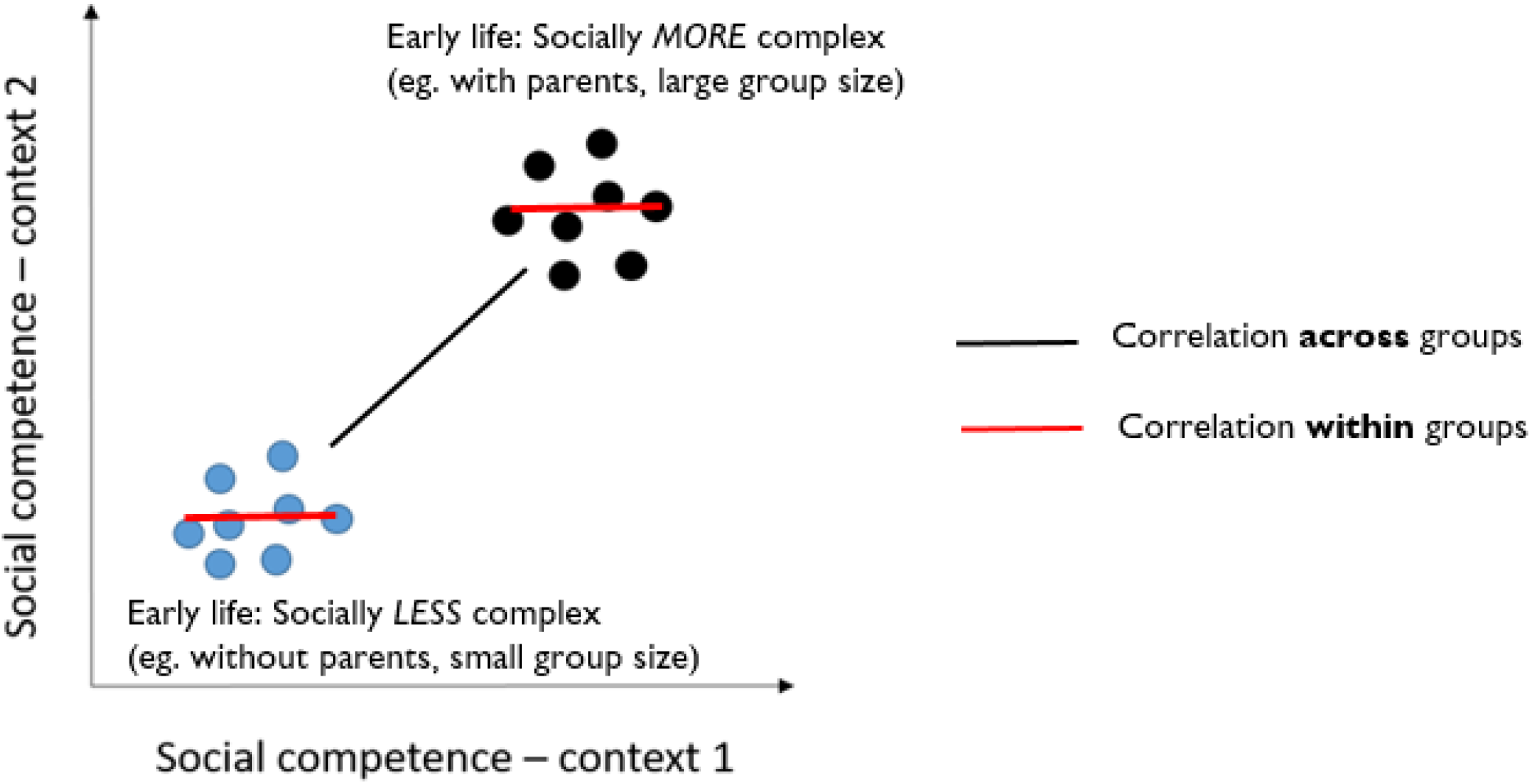
Potential reconciliation of current results with the previous developmental studies on social competence. This scheme depicts how our current results, where individuals were all reared in socially complex groups were tested repeatedly in different contexts can be reconciled with prior results on *N. pulcher* reared in different environments. Across treatment groups with socially more complex vs socially less complex early life treatments, social competence is positively correlated (black line) across contexts / roles, that is, groups that are socially competent in one context are also competent in other contexts. However, within groups, when individually tested, this relationship is largely eroded (red lines) due to interaction of social competence with consistent individual tendencies acting as a constraint for flexibility.

## Notes

https://doi.org/10.17605/OSF.IO/NPMQ2

### Competing Interest Statement

The authors have declared no competing interest.

### Summary of Updates

In this revised version, we provide a more detailed description of the dynamics of territory takeover to better justify our use of the measure of social competence. Throughout the manuscript, we have also clarified our language to avoid implying that we tested personality or stable behavioural tendencies. Instead, we suggest that the presence of stable behavioural differences in the system from previous studies may be one of the main reasons why social competence appears to be limited. We support this interpretation with additional evidence from within the study while acknowledging that, because repeated measures of individuals in both social roles were not feasible for logistical reasons, we cannot draw definitive conclusions about the stability of these behavioural differences.

