## Supplementary material for "Limits to social competence across opposing social roles in a cooperatively breeding cichlid": R code for plots and analysis - revised

Social competence - analysis


### Social competence - analysis

###### Apu Ramesh

#### 2026-08-06

In this document, we will go through the analysis of the data
collected for the project tentatively titles Social competence - what
are we measuring. The final datasets contain three csv files namely,
“fish\_id\_new.csv”, “fish\_size.csv” and “rawdata\_06112024.csv”.
fish\_id\_new.csv contains the experimental details, fish IDs and the
setup I noted, fish\_size.csv notes the sizes of focal and stimulus fish
and their elastomer tag id and rawdata\_06112024.csv is the aggregated
output obtained from BORIS after video analysis. For experimental setup
and data collection, refer to methods.docx.

#### Datasets for analysis

First, we will clean the dataset by getting rid of unwanted columns
and combining the two datasets into one analysable form. Secondary data
is also created by adding up point events or durations of state events
and rendering a final usable dataset. This step is done in R so that it
is reproducible. For all practical purposes, the output datasets are the
ones used for analysis.

For the purpose of the analysis, all the videos were checked again to
see if the intruder or both intruder and territroy owner were stressed
and exhibited stereotypic behaviours - like moving up and down the walls
of the aquarium. If this was the case after introduction, before the
intruder and territory interacted, then the trial is removed, as it is
very difficult to see when they started to interact and it inflates
latency measures.

Trials that were omitted from analysis: E20 (barrier fell), E31\_2
(barrier fell), E39 (video not recorded), E41(video not recorded),
E71(wrong sex introduced), E81(wrong focal fish), E27 (barrier fell),
E84 (video not recorded), E44 (intruder fish stressed throughout the
trial). These trials were repeated.

##### 1. Dataset to analyse relationship between variables within contexts

To analyse the relationship among variables within To or Int context,
we will use wide format of data, which is easier for correlation
analyses. For the purpose of this analysis, we can combine data from
focal as well as stimulus fish (total N = 81, Nfemale = 37, Nmale = 39,
3 - too small to sex). From the raw data, we summarise the variables to
get a secondary dataset with the following variables in columns.

“obs\_id” : Id of experiment

“date” : Date of the experiment

“time” : Time of the experiment

“focal\_id” : Id of focal fish

“focal\_status” : Status of focal fish – Territory owner (To) or
Intruder (Int)

“sex” : Sex of focal and stimulus fish

“stimulus\_id” : Id of stimulus fish

“focal\_std\_length” : Standard length (from tip of snout to base of
tail) of focal fish in mm

“stimulus\_std\_length” : Standard length (from tip of snout to base of
tail) of stimulus fish in mm

“To\_finspread\_count” : Number of fin spread events by To (Aggressive
display)

“To\_headdown\_count” : Number of head down events by To (Aggressive
display)

“To\_softtouch\_count”: Number of soft touch events by To (Submissive
display)

“To\_tailquiver\_count” : Number of tail quiver submissions shown by To
(Clear submission)

“To\_bite\_count” : Number of bites by To to Int (Overt aggression)

“To\_hook\_count” : Number of hook displays by To to Int (Submissive
display)

“To\_frontalapproach\_count” : Number of frontal approaches by To to
Int (Aggressive display)

“To\_mouthfight\_count” : Number of mouth fights between To and Int
(same as Int\_mouthfight\_count, even though it was recorded separately)
(Overt aggression)

“To\_zigzag\_count” : Number of zig-zag displays by To (Submissive
display)

“To\_sbend\_count” : Number of s-bend displays by To (Aggressive
display)

“To\_headup\_count” : Number of head-up displays by To (Submissive
display)

“To\_finspread\_duration” : Duration of fin spread event by To
(Aggressive display)

“To\_headdown\_duration” : Duration of head-down display by To
(Aggressive display)

“To\_tailquiver\_duration” : Duration of tail quiver submission (Clear
submission)

“To\_latency\_bite” : Latency of To to bite the Int

“To\_latency\_stopBite” : Latency to stop overt aggression by To to
Int

“To\_latency\_tailquiver” : Latency of To to show tail quiver
submission to Int

“To\_latency\_stoptailquiver” : Latency to stop tail quiver submission
by To to Int

“Int\_frontalapproach\_count” : Number of frontal approaches of Int to
To (Aggressive display)

“Int\_bite\_count” : Number of bites by Int to To (Overt
aggression)

“Int\_finspread\_count” : Number of fin spread events by Int to To
(Aggressive display)

“Int\_headdown\_count” : Number of head down events by Int to To
(Aggressive display)

“Int\_mouthfight\_count” : Number of mouth fights by Int and To (Overt
aggression)

“Int\_finspread\_duration” : Duration of fin spread events by Int
(Aggressive display)

“Int\_headdown\_duration” : Duration of head-down display by Int
(Aggressive display)

“Int\_latency\_bite” : Latency of Intruder to bite the To

“Int\_latency\_shelter” : Latency of Int to take over and occupy the
shelter for 5s

“Int\_latency\_stopbite” : Latency of Int to stop bites toward To

“acceptance\_score” : Acceptance score of smaller fish (prior To),
after ~3-4 hours after introduction: 0 - Evicted at the top or bottom
corner of the tank and is limited in movement, receives aggression from
the larger fish; 1- Tolerated around the shelter, free movement
possible; 2 - Fully accepted in or near shelter

“access\_shelter”: Whether the smaller fish (prior To) has access to
shelter or not after ~3-4 hours after introduction

##### 2. Dataset to analyse relationship between variables across the two contexts

To analyse the relationship among variables across To and Int
context, we will again use wide format of data, which is easier for
correlation analyses. For the purpose of this analysis, we only focus on
the focal fish, which have been subject to both the contexts (total N =
37, Nfemale = 18, Nmale = 19). We use the main\_dataset to obtain the
required columns.

##### Directly read the two datasets

##### 3. Relationship of variables of interest to the ‘success’ outcome in the two contexts

###### 3.1 Context 1: Territory owner

First, we will plot the “Binary acceptance score” (where we consider
accepted and tolerated as success with value 1 and evicted with value 0)
with the variables of interest, 1) Latency to submit, 2) Number of bites
by the TO after submitting, and 3) Number of submissions by TO. Second,
we will build separate glm models, with Binary acceptance score as the
response and each variable of interest separately, while controlling for
sex, absolute size difference between the interacting fish and the
response of the stimulus fish (here, number of bites by the INT).
Finally, we will build a multiple regression glm, with all the variables
of interest and all the controlling fixed effects, to see if there are
dependencies between our variables of interest and one stands out.

```
#We take success as fully accepted (2) and tolerated (1)

main_dataset <- main_dataset%>% 
  mutate(binary_acceptance_score = if_else(acceptance_score == 2, 1, 0))

main_dataset$binary_acceptance_score <- as.factor(main_dataset$binary_acceptance_score)

#There are some values of Inf in the submissions per received aggression in the dataset - remove these and make them NA
main_dataset[sapply(main_dataset, is.infinite)] <- NA

main_dataset$To_biteaftersubmit_persecond_count <- main_dataset$To_biteaftersubmit_count/ (1200-main_dataset$To_latency_tailquiver)

main_dataset$To_totaggaftersubmit_persecond_count <- (main_dataset$To_restaggaftersubmit_count + main_dataset$To_biteaftersubmit_count)/(1200-main_dataset$To_latency_tailquiver)


#Functions for half-violin plots

GeomSplitViolin <- ggproto("GeomSplitViolin", GeomViolin, 
                           draw_group = function(self, data, ..., draw_quantiles = NULL) {
                             data <- transform(data, xminv = x - violinwidth * (x - xmin), xmaxv = x + violinwidth * (xmax - x))
                             grp <- data[1, "group"]
                             newdata <- plyr::arrange(transform(data, x = if (grp %% 2 == 1) xminv else xmaxv), if (grp %% 2 == 1) y else -y)
                             newdata <- rbind(newdata[1, ], newdata, newdata[nrow(newdata), ], newdata[1, ])
                             newdata[c(1, nrow(newdata) - 1, nrow(newdata)), "x"] <- round(newdata[1, "x"])
                             
                             if (length(draw_quantiles) > 0 & !scales::zero_range(range(data$y))) {
                               stopifnot(all(draw_quantiles >= 0), all(draw_quantiles <=
                                                                         1))
                               quantiles <- ggplot2:::create_quantile_segment_frame(data, draw_quantiles)
                               aesthetics <- data[rep(1, nrow(quantiles)), setdiff(names(data), c("x", "y")), drop = FALSE]
                               aesthetics$alpha <- rep(1, nrow(quantiles))
                               both <- cbind(quantiles, aesthetics)
                               quantile_grob <- GeomPath$draw_panel(both, ...)
                               ggplot2:::ggname("geom_split_violin", grid::grobTree(GeomPolygon$draw_panel(newdata, ...), quantile_grob))
                             }
                             else {
                               ggplot2:::ggname("geom_split_violin", GeomPolygon$draw_panel(newdata, ...))
                             }
                           })


geom_split_violin <- function(mapping = NULL, data = NULL, stat = "ydensity", position = "identity", ..., 
                              draw_quantiles = NULL, trim = TRUE, scale = "area", na.rm = FALSE, 
                              show.legend = NA, inherit.aes = TRUE) {
  layer(data = data, mapping = mapping, stat = stat, geom = GeomSplitViolin, 
        position = position, show.legend = show.legend, inherit.aes = inherit.aes, 
        params = list(trim = trim, scale = scale, draw_quantiles = draw_quantiles, na.rm = na.rm, ...))
}


#Plots for TO

#1) Acceptance score vs Latency to submit

main_dataset$binary_acceptance_score <- factor(main_dataset$binary_acceptance_score, labels = c("Evicted", "Accepted"))


p1 <- ggplot(data=subset(main_dataset, !is.na(binary_acceptance_score)), aes(x=as.factor(binary_acceptance_score),
y=as.numeric(To_latency_tailquiver), 
fill= as.factor(binary_acceptance_score)))+
  theme_cowplot()+
  labs(y = "Latency to submit (s)", x ="Acceptance score")+
  geom_split_violin(alpha=0.5,scale = "width",  color = NA)+
  geom_boxplot( width = .3, colour = "BLACK", outliers = FALSE) +
  geom_point(position = position_jitterdodge(jitter.width = 0.15,dodge.width = 0.5), alpha = 0.3)+
  scale_y_continuous(limits = c(0, 1300))+
  guides(fill = FALSE, colour = FALSE) +
  scale_fill_manual(values =rev(RColorBrewer::brewer.pal(5,'RdBu')))+
  scale_color_manual(values =rev(RColorBrewer::brewer.pal(5,'RdBu')))+
  theme(axis.text=element_text(size=12),
        axis.title=element_text(size=17),
        strip.text.x = element_blank(),
        panel.background = element_rect(fill = 'transparent'),
        plot.background = element_rect(fill = 'transparent', color = NA)) 

#2) Acceptance score vs No.bites after submission
p2 <- ggplot(data=subset(main_dataset, !is.na(binary_acceptance_score)), aes(x=as.factor(binary_acceptance_score),
y=as.numeric(To_totaggaftersubmit_persecond_count), 
fill= as.factor(binary_acceptance_score)))+
  theme_cowplot()+
  labs(y = expression(atop("Total aggression per second", paste("after submission"))), x ="Acceptance score")+
  geom_split_violin(alpha=0.5,scale = "width",  color = NA)+
  geom_boxplot( width = .3, colour = "BLACK", outliers = FALSE) +
  geom_point(position = position_jitterdodge(jitter.width = 0.15,dodge.width = 0.5), alpha = 0.3)+
  scale_y_continuous(limits = c(0, 0.1))+
  guides(fill = FALSE, colour = FALSE) +
  scale_fill_manual(values =rev(RColorBrewer::brewer.pal(5,'RdBu')))+
  scale_color_manual(values =rev(RColorBrewer::brewer.pal(5,'RdBu')))+
  theme(axis.text=element_text(size=12),
        axis.title=element_text(size=17),
        strip.text.x = element_blank())

#3) Acceptance score vs Number of submissions per received aggression
p3 <- ggplot(data=subset(main_dataset, !is.na(binary_acceptance_score)), aes(x=as.factor(binary_acceptance_score),
y=as.numeric(To_submissionperreceivedaggression), 
fill= as.factor(binary_acceptance_score)))+
  theme_cowplot()+
  labs(y = expression(atop("Number of submissions", paste("per received bites"))), x ="Acceptance score")+
  geom_split_violin(alpha=0.5,scale = "width",  color = NA)+
  geom_boxplot( width = .3, colour = "BLACK", outliers = FALSE) +
  geom_point(position = position_jitterdodge(jitter.width = 0.15,dodge.width = 0.5), alpha = 0.3)+
  scale_y_continuous(limits = c(0, 5))+
  guides(fill = FALSE, colour = FALSE) +
  scale_fill_manual(values =rev(RColorBrewer::brewer.pal(5,'RdBu')))+
  scale_color_manual(values =rev(RColorBrewer::brewer.pal(5,'RdBu')))+
  theme(axis.text=element_text(size=12),
        axis.title=element_text(size=17),
        strip.text.x = element_blank())

plot_grid(p1, p2, p3,
          labels = c("a","b","c"), ncol = 3)
```

```
#4) Dynamics of submission plot SI 4


rawdata_1 %>%
  filter(
    subject == "Territory owner",
    behaviour %in% c("tail quiver submission", "Bite and chase")
  ) %>%
  mutate(
    submit = if_else(behaviour == "tail quiver submission", 1, 0)
  ) %>%
  ggplot(aes(x = adjusted_start,
             y = submit,
             group = obs_id)) +
  geom_point(size = 1.5, alpha = 0.6) +
  geom_smooth(
    method = "glm",
    method.args = list(family = "binomial"),
    formula = y ~ x,
    se = FALSE,
    colour = "grey70",
    linewidth = 0.8
  ) +
  scale_y_continuous(limits = c(0, 1)) +
  labs(
    x = "Time in seconds",
    y = "Probability of submission vs overt aggression",
    main = "SI.4: Submission to overt aggression over time"
  ) +
  theme_classic()
```

```
#| warning: false
#Univariate model using glm

#Binomial glm for the different variables in TO

# - After fitting, it is important to check overdispersion
# - The resulting estimates in the summary are not probabilities but log-odds ratios because we have used a logit link
# - The logit transformation takes values ranging from 0 to 1 (probabilities) and transforms them to values ranging from -Inf to +Inf. This allows us to create additive linear models without worrying about going above 1 or below 0. To get probabilities out of our model, we need to use the inverse logit. There is function for this in base R called plogis()
# - NAs in the dataset are handled explicitly in glm - we assign NA values in output as well, so that we can add the predicted values in the dataset. Otherwise it will exclude NA and lead to error when you want to plot the predicted values. Does not change the output of the glm

# Check again that there are only M and F
main_dataset$sex<-as.factor(main_dataset$sex)
main_dataset$sex[main_dataset$sex == ""] <- NA

#1) Model with Latency to tail quiver
#getting warnings about fit glm.fit: fitted probabilities numerically 0 or 1 occurred. So removing abs size diff and sex. 
to_accept1 <- glm(binary_acceptance_score ~ To_latency_tailquiver + Int_bite_count+ sex, family = binomial(link = "logit"), data = main_dataset, na.action = na.exclude)

summary(to_accept1)
```

```
## 
## Call:
## glm(formula = binary_acceptance_score ~ To_latency_tailquiver + 
##     Int_bite_count + sex, family = binomial(link = "logit"), 
##     data = main_dataset, na.action = na.exclude)
## 
## Coefficients:
##                         Estimate Std. Error z value Pr(>|z|)  
## (Intercept)            1.4131865  0.5914332   2.389   0.0169 *
## To_latency_tailquiver -0.0021286  0.0009532  -2.233   0.0255 *
## Int_bite_count        -0.0091000  0.0109148  -0.834   0.4044  
## sexM                  -0.2692987  0.5397118  -0.499   0.6178  
## ---
## Signif. codes:  0 '***' 0.001 '**' 0.01 '*' 0.05 '.' 0.1 ' ' 1
## 
## (Dispersion parameter for binomial family taken to be 1)
## 
##     Null deviance: 98.898  on 75  degrees of freedom
## Residual deviance: 92.979  on 72  degrees of freedom
##   (5 observations deleted due to missingness)
## AIC: 100.98
## 
## Number of Fisher Scoring iterations: 4
```

```
confint(to_accept1)
```

```
##                              2.5 %        97.5 %
## (Intercept)            0.303195424  2.6500716494
## To_latency_tailquiver -0.004153318 -0.0003493437
## Int_bite_count        -0.031045843  0.0127384041
## sexM                  -1.358950855  0.7760919810
```

```
var(to_accept1$residual)
```

```
## [1] 4.851629
```

```
#check overdispersion of fit - not over dispersed! 
#But the DHARma cannot handle NA, so just make a separate model without NA for it
to_accept1_noNA <- glm(binary_acceptance_score ~ To_latency_tailquiver + Int_bite_count + sex, family = binomial(link = "logit"), data = main_dataset)
summary(to_accept1_noNA)
```

```
## 
## Call:
## glm(formula = binary_acceptance_score ~ To_latency_tailquiver + 
##     Int_bite_count + sex, family = binomial(link = "logit"), 
##     data = main_dataset)
## 
## Coefficients:
##                         Estimate Std. Error z value Pr(>|z|)  
## (Intercept)            1.4131865  0.5914332   2.389   0.0169 *
## To_latency_tailquiver -0.0021286  0.0009532  -2.233   0.0255 *
## Int_bite_count        -0.0091000  0.0109148  -0.834   0.4044  
## sexM                  -0.2692987  0.5397118  -0.499   0.6178  
## ---
## Signif. codes:  0 '***' 0.001 '**' 0.01 '*' 0.05 '.' 0.1 ' ' 1
## 
## (Dispersion parameter for binomial family taken to be 1)
## 
##     Null deviance: 98.898  on 75  degrees of freedom
## Residual deviance: 92.979  on 72  degrees of freedom
##   (5 observations deleted due to missingness)
## AIC: 100.98
## 
## Number of Fisher Scoring iterations: 4
```

```
testDispersion(to_accept1_noNA)
```

```
## 
##  DHARMa nonparametric dispersion test via sd of residuals fitted vs.
##  simulated
## 
## data:  simulationOutput
## dispersion = 1.0172, p-value = 0.968
## alternative hypothesis: two.sided
```

```
#diagnostic plots
plot(to_accept1)
```

```
#2) Model with Total aggression after submission
to_accept2 <- glm(binary_acceptance_score ~ To_totaggaftersubmit_persecond_count + Int_bite_count + sex, family = binomial(link = "logit"),data = main_dataset, na.action = na.exclude)

summary(to_accept2)
```

```
## 
## Call:
## glm(formula = binary_acceptance_score ~ To_totaggaftersubmit_persecond_count + 
##     Int_bite_count + sex, family = binomial(link = "logit"), 
##     data = main_dataset, na.action = na.exclude)
## 
## Coefficients:
##                                       Estimate Std. Error z value Pr(>|z|)  
## (Intercept)                           0.809055   0.481528   1.680   0.0929 .
## To_totaggaftersubmit_persecond_count 15.597705  27.164427   0.574   0.5658  
## Int_bite_count                       -0.006859   0.010634  -0.645   0.5189  
## sexM                                 -0.044569   0.503592  -0.089   0.9295  
## ---
## Signif. codes:  0 '***' 0.001 '**' 0.01 '*' 0.05 '.' 0.1 ' ' 1
## 
## (Dispersion parameter for binomial family taken to be 1)
## 
##     Null deviance: 94.659  on 73  degrees of freedom
## Residual deviance: 93.787  on 70  degrees of freedom
##   (7 observations deleted due to missingness)
## AIC: 101.79
## 
## Number of Fisher Scoring iterations: 4
```

```
confint(to_accept2)
```

```
##                                             2.5 %      97.5 %
## (Intercept)                           -0.11897612  1.79241156
## To_totaggaftersubmit_persecond_count -28.91978936 88.93797020
## Int_bite_count                        -0.02803249  0.01466942
## sexM                                  -1.04424208  0.94355241
```

```
var(to_accept2$residual)
```

```
## [1] 4.591707
```

```
#check overdispersion of fit - not over dispersed! 
to_accept2_noNA <- glm(binary_acceptance_score ~ To_totaggaftersubmit_persecond_count + Int_bite_count + sex, family = binomial(link = "logit"),data = main_dataset)

testDispersion(to_accept2_noNA)
```

```
## 
##  DHARMa nonparametric dispersion test via sd of residuals fitted vs.
##  simulated
## 
## data:  simulationOutput
## dispersion = 1.0167, p-value = 0.944
## alternative hypothesis: two.sided
```

```
#diagnostic plots
plot(to_accept2)
```

```
#3) Model with submissions to received bites
to_accept3 <- glm(binary_acceptance_score ~ To_submissionperreceivedaggression  + Int_bite_count + sex, family = binomial(link = "logit"),data = main_dataset, na.action = na.exclude)

summary(to_accept3)
```

```
## 
## Call:
## glm(formula = binary_acceptance_score ~ To_submissionperreceivedaggression + 
##     Int_bite_count + sex, family = binomial(link = "logit"), 
##     data = main_dataset, na.action = na.exclude)
## 
## Coefficients:
##                                    Estimate Std. Error z value Pr(>|z|)  
## (Intercept)                         1.12036    0.57059   1.963   0.0496 *
## To_submissionperreceivedaggression -0.01705    0.04460  -0.382   0.7023  
## Int_bite_count                     -0.01216    0.01171  -1.039   0.2988  
## sexM                               -0.10578    0.51672  -0.205   0.8378  
## ---
## Signif. codes:  0 '***' 0.001 '**' 0.01 '*' 0.05 '.' 0.1 ' ' 1
## 
## (Dispersion parameter for binomial family taken to be 1)
## 
##     Null deviance: 91.658  on 71  degrees of freedom
## Residual deviance: 90.579  on 68  degrees of freedom
##   (9 observations deleted due to missingness)
## AIC: 98.579
## 
## Number of Fisher Scoring iterations: 4
```

```
confint(to_accept3)
```

```
##                                          2.5 %     97.5 %
## (Intercept)                         0.02569554 2.29319002
## To_submissionperreceivedaggression -0.10778780 0.08934837
## Int_bite_count                     -0.03599281 0.01110993
## sexM                               -1.13767491 0.90387000
```

```
var(to_accept3$residual)
```

```
## [1] 4.633932
```

```
#check overdispersion of fit - not over dispersed! 
to_accept3_noNA <- glm(binary_acceptance_score ~ To_submissionperreceivedaggression + Int_bite_count + sex, family = binomial,data = main_dataset)

testDispersion(to_accept3_noNA)
```

```
## 
##  DHARMa nonparametric dispersion test via sd of residuals fitted vs.
##  simulated
## 
## data:  simulationOutput
## dispersion = 1.0135, p-value = 0.944
## alternative hypothesis: two.sided
```

```
#diagnostic plots
plot(to_accept3)
```

###### 3.2 Context 2: Intruder

Similar to the context of territory owner, we analyse how the
different variables of interest are associated with the positive outcome
for Intruder. First, we will plot ‘Latency to taking over shelter’
against ‘Latency to bite’, ‘Number of bites’ and ‘Proportion of
aggressive displays’. Second, we will make univariate models with
Latency to taking over shelter as the respose variable and each of the
variable of interest as the fixed effect, while controlling for the
behaviour of the territory owner (bites by TO to INT), sex and absolute
size difference. As a third step, we also do a multiple regression model
with Latency to taking over shelter and all the behaviours of interest
as fixed effect, along with the other fixed effects.

```
#Plots for INT 

#1) Latency to shelter vs Latency to bite
i1 <- ggplot(main_dataset, aes(x = log10(Int_latency_bite),
                              y = log10(Int_latency_shelter))) +
  geom_point() +  # Add points
  theme_cowplot()+
  labs(x = "Log - Latency to bite",
       y = expression(atop("Log - Latency to", paste("take over shelter"))))+
  geom_smooth(method='lm', formula= y~x)+
  theme_bw()+
  theme(axis.text=element_text(size=12),
        axis.title=element_text(size=17),
        strip.text.x = element_blank(),
        panel.background = element_rect(fill = 'transparent'),
        plot.background = element_rect(fill = 'transparent', color = NA)) 

#2) Latency to shelter vs Number of bites
i2 <- ggplot(main_dataset, aes(x = Int_bite_count,
                              y = log10(Int_latency_shelter))) +
  geom_point() +  # Add points
  theme_cowplot()+
  labs(x = "Number of bites",
       y = expression(atop("Log - Latency to", paste("take over shelter"))))+
  geom_smooth(method='lm', formula= y~x)+
  theme_bw()+
  theme(axis.text=element_text(size=12),
        axis.title=element_text(size=17),
        strip.text.x = element_blank())

#3) Latency to shelter vs Proportion of display
i3 <- ggplot(main_dataset, aes(x = Int_prop_restagg,
                              y = log10(Int_latency_shelter))) +
  geom_point() +  # Add points
  theme_cowplot()+
  labs(x = expression(atop("Proportion of displays", paste("to total aggression"))),
       y = expression(atop("Log - Latency to", paste("take over shelter"))))+
  geom_smooth(method='lm', formula= y~x)+
  theme_bw()+
  theme(axis.text=element_text(size=12),
        axis.title=element_text(size=17),
        strip.text.x = element_blank())


plot_grid(i1, i2, i3,
          labels = c("a","b","c"), ncol = 3)
```

```
title <- ggdraw() +
  draw_label(
    "Figure 2",
    fontface = "bold",
    x = 0, hjust = 0,
    size = 16
  )

panel <- plot_grid(p1, p2, p3,i1, i2, i3,
          labels = c("a","b","c", "d", "e", "f"),
          ncol = 3)

final_plot <- plot_grid(
  title,
  panel,
  ncol = 1,
  rel_heights = c(0.08, 1)
)

final_plot
```

```
# Supplementary plot for INT aggression dynamics
# Intruder attacks with time, centred around latency to takeover shelter

takeover <- rawdata_1 %>%
  filter(behaviour == "latency for intruder to occupy the shelter for the first time for more than 5 seconds") %>%
  dplyr::select(obs_id, takeover_time = adjusted_start)

bites <- rawdata_1 %>%
  dplyr::filter(behaviour == "Bite and chase") %>%
  dplyr::left_join(takeover, by = "obs_id") %>%
  dplyr::mutate(
    rel_time = adjusted_start - takeover_time
  )
bites <- bites %>%
    filter(rel_time >= -1200,
           rel_time <= 1200)

#create 10 second bite bins
bite_counts <- bites %>%
    mutate(time10 = floor(rel_time/10)*10) %>%
    count(obs_id, time10)

# fill in missing time bins
library(tidyr)

all_times <- seq(-1200,1200,10)

bite_counts <- bite_counts %>%
    complete(
        obs_id,
        time10 = all_times,
        fill=list(n=0)
    )

# fit gam model
library(mgcv)
bite_counts$obs_id <- factor(bite_counts$obs_id)
m <- gam(
    n ~
      s(time10, k=20) +
      s(obs_id, bs="re"),
    family=nb(),
    data=bite_counts,
    method="REML"
)

# Get prediction from gam model

newdat <- data.frame(
  time10 = seq(
    min(bite_counts$time10, na.rm = TRUE),
    max(bite_counts$time10, na.rm = TRUE),
    length.out = 300
  ),
  obs_id = factor(
    bite_counts$obs_id[1],
    levels = levels(bite_counts$obs_id)
  )
)

X <- predict(
  m,
  newdata = newdat,
  type = "lpmatrix",
  exclude = "s(obs_id)"
)

beta <- coef(m)
V <- vcov(m)

fit <- X %*% beta
se  <- sqrt(diag(X %*% V %*% t(X)))

newdat$fit <- exp(fit)
newdat$upper <- exp(fit + 2 * se)
newdat$lower <- exp(fit - 2 * se)

plot(newdat$time10, newdat$fit, type = "l",
     lwd = 2,
     xlab = "Time (seconds)",
     ylab = "Predicted overt attacks",
     main = "SI.5: Overt attack intensity over time",
     ylim = c(0, 0.6)
     )
lines(newdat$time10, newdat$upper, lty = 2)
lines(newdat$time10, newdat$lower, lty = 2)
```

```
#1) Latency to take over shelter vs Latency to bite

int_shel1 <- glm(Int_latency_shelter~ Int_latency_bite + To_bite_count + sex, family = gaussian,data = main_dataset, na.action = na.exclude)

summary(int_shel1)
```

```
## 
## Call:
## glm(formula = Int_latency_shelter ~ Int_latency_bite + To_bite_count + 
##     sex, family = gaussian, data = main_dataset, na.action = na.exclude)
## 
## Coefficients:
##                  Estimate Std. Error t value Pr(>|t|)    
## (Intercept)       53.0824    48.1102   1.103    0.274    
## Int_latency_bite   0.6330     0.0923   6.859 2.01e-09 ***
## To_bite_count      8.8360     1.5704   5.627 3.30e-07 ***
## sexM              89.1330    56.3976   1.580    0.118    
## ---
## Signif. codes:  0 '***' 0.001 '**' 0.01 '*' 0.05 '.' 0.1 ' ' 1
## 
## (Dispersion parameter for gaussian family taken to be 58485.43)
## 
##     Null deviance: 9996480  on 75  degrees of freedom
## Residual deviance: 4210951  on 72  degrees of freedom
##   (5 observations deleted due to missingness)
## AIC: 1055.8
## 
## Number of Fisher Scoring iterations: 2
```

```
confint(int_shel1)
```

```
##                       2.5 %      97.5 %
## (Intercept)      -41.211930 147.3766469
## Int_latency_bite   0.452141   0.8139489
## To_bite_count      5.758086  11.9139563
## sexM             -21.404218 199.6702425
```

```
var(int_shel1$residual)
```

```
## [1] 56146.02
```

```
plot(int_shel1)
```

```
hist(resid(int_shel1)) #Long tails on one side but otherwise looks fine
```

```
#2) Latency to shelter vs Number of bites
int_shel2_2 <- glm(Int_latency_shelter~ Int_bite_count + To_bite_count + sex, family = gaussian,data = main_dataset, na.action = na.exclude)

summary(int_shel2_2)
```

```
## 
## Call:
## glm(formula = Int_latency_shelter ~ Int_bite_count + To_bite_count + 
##     sex, family = gaussian, data = main_dataset, na.action = na.exclude)
## 
## Coefficients:
##                Estimate Std. Error t value Pr(>|t|)    
## (Intercept)     291.879     66.578   4.384 3.90e-05 ***
## Int_bite_count   -4.062      1.510  -2.690  0.00888 ** 
## To_bite_count    11.494      1.890   6.082 5.17e-08 ***
## sexM             41.262     69.451   0.594  0.55429    
## ---
## Signif. codes:  0 '***' 0.001 '**' 0.01 '*' 0.05 '.' 0.1 ' ' 1
## 
## (Dispersion parameter for gaussian family taken to be 87866.53)
## 
##     Null deviance: 9996480  on 75  degrees of freedom
## Residual deviance: 6326390  on 72  degrees of freedom
##   (5 observations deleted due to missingness)
## AIC: 1086.7
## 
## Number of Fisher Scoring iterations: 2
```

```
confint(int_shel2_2)
```

```
##                     2.5 %     97.5 %
## (Intercept)    161.388682 422.368345
## Int_bite_count  -7.021593  -1.102152
## To_bite_count    7.790336  15.198525
## sexM           -94.859650 177.384208
```

```
var(int_shel2_2$residual)
```

```
## [1] 84351.87
```

```
hist(resid(int_shel2_2)) # Again, long tails on one side but otherwise looks fine
```

```
# testDispersion(int_shel2_2) - can only be tested when you dont have  na.action = na.exclude in glm specifications
#3) Latency to shelter vs Proportion of displays to total aggression
int_shel3 <- glm(Int_latency_shelter ~ Int_prop_restagg + To_bite_count + sex, family = gaussian,data = main_dataset, na.action = na.exclude)

summary(int_shel3)
```

```
## 
## Call:
## glm(formula = Int_latency_shelter ~ Int_prop_restagg + To_bite_count + 
##     sex, family = gaussian, data = main_dataset, na.action = na.exclude)
## 
## Coefficients:
##                  Estimate Std. Error t value Pr(>|t|)    
## (Intercept)       -13.378     70.778  -0.189 0.850622    
## Int_prop_restagg  449.460    121.692   3.693 0.000431 ***
## To_bite_count      11.181      1.766   6.332 1.92e-08 ***
## sexM              115.716     65.763   1.760 0.082785 .  
## ---
## Signif. codes:  0 '***' 0.001 '**' 0.01 '*' 0.05 '.' 0.1 ' ' 1
## 
## (Dispersion parameter for gaussian family taken to be 77169.52)
## 
##     Null deviance: 9643037  on 74  degrees of freedom
## Residual deviance: 5479036  on 71  degrees of freedom
##   (6 observations deleted due to missingness)
## AIC: 1062.8
## 
## Number of Fisher Scoring iterations: 2
```

```
confint(int_shel3)
```

```
##                        2.5 %    97.5 %
## (Intercept)      -152.100624 125.34470
## Int_prop_restagg  210.947845 687.97267
## To_bite_count       7.720409  14.64232
## sexM              -13.177194 244.60889
```

```
var(int_shel3$residual)
```

```
## [1] 74041.02
```

```
plot(int_shel3)
```

```
hist(resid(int_shel3))
```

```
# testDispersion(int_shel3)
#4) Latency to shelter vs all
int_shel4 <- glm(Int_latency_shelter ~ Int_latency_bite + Int_bite_count + Int_prop_restagg + To_bite_count + sex + abs_sizediff, family = gaussian,data = main_dataset, na.action = na.exclude)

summary(int_shel4)
```

```
## 
## Call:
## glm(formula = Int_latency_shelter ~ Int_latency_bite + Int_bite_count + 
##     Int_prop_restagg + To_bite_count + sex + abs_sizediff, family = gaussian, 
##     data = main_dataset, na.action = na.exclude)
## 
## Coefficients:
##                  Estimate Std. Error t value Pr(>|t|)    
## (Intercept)      334.1793   218.4481   1.530   0.1310    
## Int_latency_bite   0.6407     0.1289   4.971 5.26e-06 ***
## Int_bite_count     0.7428     1.5401   0.482   0.6312    
## Int_prop_restagg  68.3238   144.6711   0.472   0.6383    
## To_bite_count      8.2315     1.7186   4.790 1.03e-05 ***
## sexM             131.4426    63.2988   2.077   0.0419 *  
## abs_sizediff     -37.7555    20.8946  -1.807   0.0755 .  
## ---
## Signif. codes:  0 '***' 0.001 '**' 0.01 '*' 0.05 '.' 0.1 ' ' 1
## 
## (Dispersion parameter for gaussian family taken to be 60389.12)
## 
##     Null deviance: 9306599  on 70  degrees of freedom
## Residual deviance: 3864904  on 64  degrees of freedom
##   (10 observations deleted due to missingness)
## AIC: 991.73
## 
## Number of Fisher Scoring iterations: 2
```

```
plot(int_shel4)
```

```
hist(resid(int_shel4))
```

```
# testDispersion(int_shel4)
```

##### 4. Relationship between the two outcome variables

```
# Acceptance score vs Latency to take over shelter
focal_to_int<- focal_to_int%>% 
  mutate(binary_acceptance_score = if_else(acceptance_score == 2, 1, 0))
  
focal_to_int$binary_acceptance_score <- factor(focal_to_int$binary_acceptance_score, labels = c("Evicted", "Accepted"))

focal_to_int$To_biteaftersubmit_persecond_count <- focal_to_int$To_biteaftersubmit_count/ (1200-focal_to_int$To_latency_tailquiver)

focal_to_int$To_totaggaftersubmit_persecond_count <- (focal_to_int$To_restaggaftersubmit_count + focal_to_int$To_biteaftersubmit_count)/(1200-focal_to_int$To_latency_tailquiver)


plot1 <- ggplot(data=subset(focal_to_int, !is.na(binary_acceptance_score)), aes(x=as.factor(binary_acceptance_score),
y=as.numeric(Int_latency_shelter), 
fill= as.factor(binary_acceptance_score)))+
  theme_cowplot()+
  labs(y = "INT: Latency to take over shelter (s)", x ="TO: Acceptance score",
       title = "SI.6 : Relationship between the outcomes")+
  geom_split_violin(alpha=0.5,scale = "width",  color = NA)+
  geom_boxplot( width = .3, colour = "BLACK", outliers = FALSE) +
  geom_point( position = position_jitterdodge(jitter.width = 0.15,dodge.width = 0.5), alpha = 0.3)+
  scale_y_continuous(limits = c(0, 1300))+
  guides(fill = FALSE, colour = FALSE) +
  scale_fill_manual(values =rev(RColorBrewer::brewer.pal(5,'RdBu')))+
  scale_color_manual(values =rev(RColorBrewer::brewer.pal(5,'RdBu')))+
  theme(axis.text=element_text(size=12),
        axis.title=element_text(size=17),
        strip.text.x = element_blank(),
        panel.background = element_rect(fill = 'transparent'),
        plot.background = element_rect(fill = 'transparent', color = NA))

plot1
```

```
#non parametric wilco test to see if latency to take over shelter is different between accepted and evicted fish

accepted_fish <- subset(focal_to_int, binary_acceptance_score == "Accepted")

evicted_fish <- subset(focal_to_int, binary_acceptance_score == "Evicted")

wilcox.test(accepted_fish$Int_latency_shelter, evicted_fish$Int_latency_shelter)
```

```
## 
##  Wilcoxon rank sum test with continuity correction
## 
## data:  accepted_fish$Int_latency_shelter and evicted_fish$Int_latency_shelter
## W = 190.5, p-value = 0.2794
## alternative hypothesis: true location shift is not equal to 0
```

##### 4. Relationship among competent behaviours across roles

###### 4.1 Raw correlations

```
#custom function for spearman correlation matrix
chartcorr_spearman<-function (R, histogram = TRUE, method =
                                "spearman", ...)
{
  x = checkData(R, method = "matrix")
  if (missing(method))
    method = method[1]
  panel.cor <- function(x, y, digits = 2, prefix = "", use = "pairwise.complete.obs",
                        method = "spearman", cex.cor, ...) {
    usr <- par("usr")
    on.exit(par(usr))
    par(usr = c(0, 1, 0, 1))
    r <- cor(x, y, use = use, method = method)
    txt <- format(c(r, 0.123456789), digits = digits)[1]
    txt <- paste(prefix, txt, sep = "")
    if (missing(cex.cor))
      cex <- 0.8/strwidth(txt)
    test <- cor.test(as.numeric(x), as.numeric(y), method = method)
    Signif <- symnum(test$p.value, corr = FALSE, na = FALSE,
                     cutpoints = c(0, 0.001, 0.01, 0.05, 0.1, 1), symbols = c("***",
                                                                              "**", "*", ".", " "))
    text(0.5, 0.5, txt, cex = cex * (abs(r) + 0.3)/1.3)
    text(0.8, 0.8, Signif, cex = cex, col = 2)
  }
  f <- function(t) {
    dnorm(t, mean = mean(x), sd = sd.xts(x))
  }
  dotargs <- list(...)
  dotargs$method <- NULL
  rm(method)
  hist.panel = function(x, ... = NULL) {
    par(new = TRUE)
    hist(x, col = "light gray", probability = TRUE, axes = FALSE,
         main = "", breaks = "FD")
    lines(density(x, na.rm = TRUE), col = "red", lwd = 1)
    rug(x)
  }
  if (histogram)
    pairs(x, gap = 0, lower.panel = panel.smooth, upper.panel = panel.cor,
          diag.panel = hist.panel)
  else pairs(x, gap = 0, lower.panel = panel.smooth, upper.panel = panel.cor)
}

#Raw correlations within and across contexts. 


cor_plot <- subset(focal_to_int, select = c(To_latency_tailquiver,
                                            To_totaggaftersubmit_persecond_count,
                                            To_submissionperreceivedaggression,
                                            Int_latency_bite,
                                            Int_bite_count,
                                            Int_prop_restagg
                                            ))
cor_plot[sapply(cor_plot, is.infinite)] <- NA
chartcorr_spearman(cor_plot)
title("SI.7: Spearman correlation matrix", 
      outer = TRUE, 
      line = 1, 
      cex.main = 1.5, 
      font.main = 2)
```

##### 5. Multivariate model to get the residual correlations, after transforming the data to gaussian using BRMS

```
#1) Scale the variables: Latency data and proportion data are assumed to be normally distributed (note that the data need not be normally distributed but the residuals, after fitting the model). Then we log -transform the count data +1. +1 is added to avoid log(0). In addition, we scale all variables (standardize) - to get values between 0 and 1

focal_to_int$To_scaled_latency_tailquiver <- as.data.frame(scale(focal_to_int$To_latency_tailquiver))[[1]]

focal_to_int$To_scaled_tailquiver_count <- as.data.frame(scale(log(focal_to_int$To_tailquiver_count+1)))[[1]]

focal_to_int$To_scaled_bitepersecond_aftersubmit_count <- as.data.frame(scale(focal_to_int$To_totaggaftersubmit_persecond_count))[[1]]

focal_to_int$TO_scaled_int_bite <- as.data.frame(scale(log(focal_to_int$TO_Int_bite_count+1)))[[1]]


focal_to_int$Int_scaled_latency_bite <- as.data.frame(scale(focal_to_int$Int_latency_bite))[[1]]

focal_to_int$Int_scaled_bite_count <- as.data.frame(scale(log(focal_to_int$Int_bite_count+1)))[[1]]

focal_to_int$Int_To_scaled_bite_count <- as.data.frame(scale(log(focal_to_int$INT_To_bite_count +1)))[[1]]


#2) Verbose multivariate analysis, controlling for different variables
#for different responses, with scaled variables, against the stimulus response


#a) To latency submit
bf_To_submit_latency1 <- bf( To_scaled_latency_tailquiver ~ TO_scaled_int_bite + sex) + gaussian()

#b) To submission count
bf_To_submit_count1 <- bf(To_scaled_tailquiver_count ~ TO_scaled_int_bite+ sex) + gaussian()

#c) To bites after submission count
bf_To_biteaftersubmit_count1 <- bf(To_scaled_bitepersecond_aftersubmit_count ~ TO_scaled_int_bite+ sex) + gaussian()

#d) Int latency bite
bf_Int_bite_latency1 <- bf(Int_scaled_latency_bite ~ Int_To_scaled_bite_count+ sex) + gaussian()

#e) Int bite count
bf_Int_bite_count1 <- bf(Int_scaled_bite_count ~ Int_To_scaled_bite_count + sex) + gaussian()

#f) Int prop rest agg - prior for proportion should be bounded between 0 and 1
bf_Int_restgg1 <- bf(Int_prop_restagg ~ Int_To_scaled_bite_count + sex) + gaussian()

#g) Final model is the combination of all the models
bf_total1 <- bf_To_submit_latency1 +
  bf_To_submit_count1 +
  bf_To_biteaftersubmit_count1 +
  bf_Int_bite_latency1 +
  bf_Int_bite_count1 +
  bf_Int_restgg1 +
  set_rescor(rescor = TRUE)

#3) Check the default priors and set good priors, especially for slopes
get_prior(formula = bf_total1, data = focal_to_int )
```

```
#4) Fit and run the brms model
fit1 <- brms::brm(bf_total1,
            data = focal_to_int,
            warmup=1000,
            iter=5000,
            chains=4,
            cores=2,
            backend = "cmdstanr",
            control = list(adapt_delta = 0.90,
                           max_treedepth = 10),
            file="mv_mode_1_")

#5) Adding information criterion to models to use for selection

fit1_1 <- add_criterion(fit1, "loo")

#6)
summary(fit1_1)
```

```
##  Family: MV(gaussian, gaussian, gaussian, gaussian, gaussian, gaussian) 
##   Links: mu = identity
##          mu = identity
##          mu = identity
##          mu = identity
##          mu = identity
##          mu = identity 
## Formula: To_scaled_latency_tailquiver ~ TO_scaled_int_bite + sex 
##          To_scaled_tailquiver_count ~ TO_scaled_int_bite + sex 
##          To_scaled_bitepersecond_aftersubmit_count ~ TO_scaled_int_bite + sex 
##          Int_scaled_latency_bite ~ Int_To_scaled_bite_count + sex 
##          Int_scaled_bite_count ~ Int_To_scaled_bite_count + sex 
##          Int_prop_restagg ~ Int_To_scaled_bite_count + sex 
##    Data: focal_to_int (Number of observations: 36) 
##   Draws: 4 chains, each with iter = 5000; warmup = 1000; thin = 1;
##          total post-warmup draws = 16000
## 
## Regression Coefficients:
##                                                          Estimate Est.Error
## Toscaledlatencytailquiver_Intercept                         -0.62      0.90
## Toscaledtailquivercount_Intercept                            0.81      0.99
## Toscaledbitepersecondaftersubmitcount_Intercept             -0.34      1.20
## Intscaledlatencybite_Intercept                              -0.38      1.01
## Intscaledbitecount_Intercept                                 0.47      0.96
## Intproprestagg_Intercept                                     0.45      0.28
## Toscaledlatencytailquiver_TO_scaled_int_bite                -0.26      0.15
## Toscaledlatencytailquiver_sexF                               0.88      0.93
## Toscaledlatencytailquiver_sexM                               0.34      0.92
## Toscaledtailquivercount_TO_scaled_int_bite                   0.23      0.16
## Toscaledtailquivercount_sexF                                -0.71      1.02
## Toscaledtailquivercount_sexM                                -0.88      1.02
## Toscaledbitepersecondaftersubmitcount_TO_scaled_int_bite    -0.03      0.19
## Toscaledbitepersecondaftersubmitcount_sexF                   0.19      1.24
## Toscaledbitepersecondaftersubmitcount_sexM                   0.52      1.23
## Intscaledlatencybite_Int_To_scaled_bite_count                0.33      0.19
## Intscaledlatencybite_sexF                                    0.08      1.05
## Intscaledlatencybite_sexM                                    0.55      1.03
## Intscaledbitecount_Int_To_scaled_bite_count                  0.00      0.18
## Intscaledbitecount_sexF                                     -0.01      1.00
## Intscaledbitecount_sexM                                     -0.81      0.98
## Intproprestagg_Int_To_scaled_bite_count                      0.01      0.05
## Intproprestagg_sexF                                         -0.13      0.30
## Intproprestagg_sexM                                         -0.01      0.29
##                                                          l-95% CI u-95% CI Rhat
## Toscaledlatencytailquiver_Intercept                         -2.38     1.17 1.00
## Toscaledtailquivercount_Intercept                           -1.14     2.77 1.00
## Toscaledbitepersecondaftersubmitcount_Intercept             -2.69     2.02 1.00
## Intscaledlatencybite_Intercept                              -2.35     1.64 1.00
## Intscaledbitecount_Intercept                                -1.45     2.33 1.00
## Intproprestagg_Intercept                                    -0.11     1.01 1.00
## Toscaledlatencytailquiver_TO_scaled_int_bite                -0.55     0.03 1.00
## Toscaledlatencytailquiver_sexF                              -0.94     2.70 1.00
## Toscaledlatencytailquiver_sexM                              -1.49     2.15 1.00
## Toscaledtailquivercount_TO_scaled_int_bite                  -0.09     0.54 1.00
## Toscaledtailquivercount_sexF                                -2.75     1.31 1.00
## Toscaledtailquivercount_sexM                                -2.89     1.13 1.00
## Toscaledbitepersecondaftersubmitcount_TO_scaled_int_bite    -0.41     0.34 1.00
## Toscaledbitepersecondaftersubmitcount_sexF                  -2.25     2.59 1.00
## Toscaledbitepersecondaftersubmitcount_sexM                  -1.89     2.92 1.00
## Intscaledlatencybite_Int_To_scaled_bite_count               -0.03     0.71 1.00
## Intscaledlatencybite_sexF                                   -2.03     2.13 1.00
## Intscaledlatencybite_sexM                                   -1.51     2.58 1.00
## Intscaledbitecount_Int_To_scaled_bite_count                 -0.34     0.36 1.00
## Intscaledbitecount_sexF                                     -1.94     1.99 1.00
## Intscaledbitecount_sexM                                     -2.73     1.17 1.00
## Intproprestagg_Int_To_scaled_bite_count                     -0.10     0.12 1.00
## Intproprestagg_sexF                                         -0.72     0.46 1.00
## Intproprestagg_sexM                                         -0.59     0.57 1.00
##                                                          Bulk_ESS Tail_ESS
## Toscaledlatencytailquiver_Intercept                          8336     9774
## Toscaledtailquivercount_Intercept                            8391     9259
## Toscaledbitepersecondaftersubmitcount_Intercept              9442    10289
## Intscaledlatencybite_Intercept                               7909     9062
## Intscaledbitecount_Intercept                                 6753     8829
## Intproprestagg_Intercept                                     6945     9186
## Toscaledlatencytailquiver_TO_scaled_int_bite                13996    11745
## Toscaledlatencytailquiver_sexF                               8377    10020
## Toscaledlatencytailquiver_sexM                               8504     9619
## Toscaledtailquivercount_TO_scaled_int_bite                  14221    11898
## Toscaledtailquivercount_sexF                                 8323     9196
## Toscaledtailquivercount_sexM                                 8554     9874
## Toscaledbitepersecondaftersubmitcount_TO_scaled_int_bite    15310    11451
## Toscaledbitepersecondaftersubmitcount_sexF                   9395    10031
## Toscaledbitepersecondaftersubmitcount_sexM                   9631    10238
## Intscaledlatencybite_Int_To_scaled_bite_count               10448    11389
## Intscaledlatencybite_sexF                                    7962     9398
## Intscaledlatencybite_sexM                                    7940     9971
## Intscaledbitecount_Int_To_scaled_bite_count                  9606     9743
## Intscaledbitecount_sexF                                      6830     9020
## Intscaledbitecount_sexM                                      6828     9342
## Intproprestagg_Int_To_scaled_bite_count                     10434    11166
## Intproprestagg_sexF                                          7003     8781
## Intproprestagg_sexM                                          7041     9508
## 
## Further Distributional Parameters:
##                                             Estimate Est.Error l-95% CI
## sigma_Toscaledlatencytailquiver                 0.90      0.12     0.70
## sigma_Toscaledtailquivercount                   0.98      0.13     0.76
## sigma_Toscaledbitepersecondaftersubmitcount     1.18      0.16     0.92
## sigma_Intscaledlatencybite                      1.00      0.13     0.78
## sigma_Intscaledbitecount                        0.95      0.12     0.74
## sigma_Intproprestagg                            0.28      0.04     0.22
##                                             u-95% CI Rhat Bulk_ESS Tail_ESS
## sigma_Toscaledlatencytailquiver                 1.16 1.00    13639    12005
## sigma_Toscaledtailquivercount                   1.28 1.00    13426    11224
## sigma_Toscaledbitepersecondaftersubmitcount     1.55 1.00    14530    11621
## sigma_Intscaledlatencybite                      1.29 1.00    12508    11674
## sigma_Intscaledbitecount                        1.22 1.00    11160    11195
## sigma_Intproprestagg                            0.37 1.00    12422    11776
## 
## Residual Correlations: 
##                                                                         Estimate
## rescor(Toscaledlatencytailquiver,Toscaledtailquivercount)                  -0.32
## rescor(Toscaledlatencytailquiver,Toscaledbitepersecondaftersubmitcount)    -0.15
## rescor(Toscaledtailquivercount,Toscaledbitepersecondaftersubmitcount)      -0.19
## rescor(Toscaledlatencytailquiver,Intscaledlatencybite)                     -0.15
## rescor(Toscaledtailquivercount,Intscaledlatencybite)                        0.19
## rescor(Toscaledbitepersecondaftersubmitcount,Intscaledlatencybite)          0.04
## rescor(Toscaledlatencytailquiver,Intscaledbitecount)                        0.17
## rescor(Toscaledtailquivercount,Intscaledbitecount)                         -0.15
## rescor(Toscaledbitepersecondaftersubmitcount,Intscaledbitecount)            0.16
## rescor(Intscaledlatencybite,Intscaledbitecount)                            -0.53
## rescor(Toscaledlatencytailquiver,Intproprestagg)                           -0.18
## rescor(Toscaledtailquivercount,Intproprestagg)                              0.12
## rescor(Toscaledbitepersecondaftersubmitcount,Intproprestagg)                0.04
## rescor(Intscaledlatencybite,Intproprestagg)                                 0.36
## rescor(Intscaledbitecount,Intproprestagg)                                  -0.57
##                                                                         Est.Error
## rescor(Toscaledlatencytailquiver,Toscaledtailquivercount)                    0.15
## rescor(Toscaledlatencytailquiver,Toscaledbitepersecondaftersubmitcount)      0.16
## rescor(Toscaledtailquivercount,Toscaledbitepersecondaftersubmitcount)        0.16
## rescor(Toscaledlatencytailquiver,Intscaledlatencybite)                       0.17
## rescor(Toscaledtailquivercount,Intscaledlatencybite)                         0.16
## rescor(Toscaledbitepersecondaftersubmitcount,Intscaledlatencybite)           0.16
## rescor(Toscaledlatencytailquiver,Intscaledbitecount)                         0.17
## rescor(Toscaledtailquivercount,Intscaledbitecount)                           0.17
## rescor(Toscaledbitepersecondaftersubmitcount,Intscaledbitecount)             0.16
## rescor(Intscaledlatencybite,Intscaledbitecount)                              0.12
## rescor(Toscaledlatencytailquiver,Intproprestagg)                             0.17
## rescor(Toscaledtailquivercount,Intproprestagg)                               0.17
## rescor(Toscaledbitepersecondaftersubmitcount,Intproprestagg)                 0.16
## rescor(Intscaledlatencybite,Intproprestagg)                                  0.15
## rescor(Intscaledbitecount,Intproprestagg)                                    0.12
##                                                                         l-95% CI
## rescor(Toscaledlatencytailquiver,Toscaledtailquivercount)                  -0.59
## rescor(Toscaledlatencytailquiver,Toscaledbitepersecondaftersubmitcount)    -0.45
## rescor(Toscaledtailquivercount,Toscaledbitepersecondaftersubmitcount)      -0.49
## rescor(Toscaledlatencytailquiver,Intscaledlatencybite)                     -0.47
## rescor(Toscaledtailquivercount,Intscaledlatencybite)                       -0.14
## rescor(Toscaledbitepersecondaftersubmitcount,Intscaledlatencybite)         -0.28
## rescor(Toscaledlatencytailquiver,Intscaledbitecount)                       -0.17
## rescor(Toscaledtailquivercount,Intscaledbitecount)                         -0.45
## rescor(Toscaledbitepersecondaftersubmitcount,Intscaledbitecount)           -0.16
## rescor(Intscaledlatencybite,Intscaledbitecount)                            -0.74
## rescor(Toscaledlatencytailquiver,Intproprestagg)                           -0.50
## rescor(Toscaledtailquivercount,Intproprestagg)                             -0.21
## rescor(Toscaledbitepersecondaftersubmitcount,Intproprestagg)               -0.29
## rescor(Intscaledlatencybite,Intproprestagg)                                 0.05
## rescor(Intscaledbitecount,Intproprestagg)                                  -0.77
##                                                                         u-95% CI
## rescor(Toscaledlatencytailquiver,Toscaledtailquivercount)                  -0.00
## rescor(Toscaledlatencytailquiver,Toscaledbitepersecondaftersubmitcount)     0.19
## rescor(Toscaledtailquivercount,Toscaledbitepersecondaftersubmitcount)       0.14
## rescor(Toscaledlatencytailquiver,Intscaledlatencybite)                      0.19
## rescor(Toscaledtailquivercount,Intscaledlatencybite)                        0.49
## rescor(Toscaledbitepersecondaftersubmitcount,Intscaledlatencybite)          0.36
## rescor(Toscaledlatencytailquiver,Intscaledbitecount)                        0.48
## rescor(Toscaledtailquivercount,Intscaledbitecount)                          0.19
## rescor(Toscaledbitepersecondaftersubmitcount,Intscaledbitecount)            0.46
## rescor(Intscaledlatencybite,Intscaledbitecount)                            -0.25
## rescor(Toscaledlatencytailquiver,Intproprestagg)                            0.17
## rescor(Toscaledtailquivercount,Intproprestagg)                              0.44
## rescor(Toscaledbitepersecondaftersubmitcount,Intproprestagg)                0.35
## rescor(Intscaledlatencybite,Intproprestagg)                                 0.62
## rescor(Intscaledbitecount,Intproprestagg)                                  -0.31
##                                                                         Rhat
## rescor(Toscaledlatencytailquiver,Toscaledtailquivercount)               1.00
## rescor(Toscaledlatencytailquiver,Toscaledbitepersecondaftersubmitcount) 1.00
## rescor(Toscaledtailquivercount,Toscaledbitepersecondaftersubmitcount)   1.00
## rescor(Toscaledlatencytailquiver,Intscaledlatencybite)                  1.00
## rescor(Toscaledtailquivercount,Intscaledlatencybite)                    1.00
## rescor(Toscaledbitepersecondaftersubmitcount,Intscaledlatencybite)      1.00
## rescor(Toscaledlatencytailquiver,Intscaledbitecount)                    1.00
## rescor(Toscaledtailquivercount,Intscaledbitecount)                      1.00
## rescor(Toscaledbitepersecondaftersubmitcount,Intscaledbitecount)        1.00
## rescor(Intscaledlatencybite,Intscaledbitecount)                         1.00
## rescor(Toscaledlatencytailquiver,Intproprestagg)                        1.00
## rescor(Toscaledtailquivercount,Intproprestagg)                          1.00
## rescor(Toscaledbitepersecondaftersubmitcount,Intproprestagg)            1.00
## rescor(Intscaledlatencybite,Intproprestagg)                             1.00
## rescor(Intscaledbitecount,Intproprestagg)                               1.00
##                                                                         Bulk_ESS
## rescor(Toscaledlatencytailquiver,Toscaledtailquivercount)                  13835
## rescor(Toscaledlatencytailquiver,Toscaledbitepersecondaftersubmitcount)    14023
## rescor(Toscaledtailquivercount,Toscaledbitepersecondaftersubmitcount)      13836
## rescor(Toscaledlatencytailquiver,Intscaledlatencybite)                     11827
## rescor(Toscaledtailquivercount,Intscaledlatencybite)                       10589
## rescor(Toscaledbitepersecondaftersubmitcount,Intscaledlatencybite)         11474
## rescor(Toscaledlatencytailquiver,Intscaledbitecount)                        9837
## rescor(Toscaledtailquivercount,Intscaledbitecount)                         10891
## rescor(Toscaledbitepersecondaftersubmitcount,Intscaledbitecount)           10566
## rescor(Intscaledlatencybite,Intscaledbitecount)                             9634
## rescor(Toscaledlatencytailquiver,Intproprestagg)                           11331
## rescor(Toscaledtailquivercount,Intproprestagg)                             11559
## rescor(Toscaledbitepersecondaftersubmitcount,Intproprestagg)               11793
## rescor(Intscaledlatencybite,Intproprestagg)                                 9654
## rescor(Intscaledbitecount,Intproprestagg)                                   9742
##                                                                         Tail_ESS
## rescor(Toscaledlatencytailquiver,Toscaledtailquivercount)                  11423
## rescor(Toscaledlatencytailquiver,Toscaledbitepersecondaftersubmitcount)    11289
## rescor(Toscaledtailquivercount,Toscaledbitepersecondaftersubmitcount)      11386
## rescor(Toscaledlatencytailquiver,Intscaledlatencybite)                     12313
## rescor(Toscaledtailquivercount,Intscaledlatencybite)                       10878
## rescor(Toscaledbitepersecondaftersubmitcount,Intscaledlatencybite)         11876
## rescor(Toscaledlatencytailquiver,Intscaledbitecount)                        9972
## rescor(Toscaledtailquivercount,Intscaledbitecount)                         11691
## rescor(Toscaledbitepersecondaftersubmitcount,Intscaledbitecount)           11628
## rescor(Intscaledlatencybite,Intscaledbitecount)                            11693
## rescor(Toscaledlatencytailquiver,Intproprestagg)                           12074
## rescor(Toscaledtailquivercount,Intproprestagg)                             12219
## rescor(Toscaledbitepersecondaftersubmitcount,Intproprestagg)               12642
## rescor(Intscaledlatencybite,Intproprestagg)                                11513
## rescor(Intscaledbitecount,Intproprestagg)                                  11457
## 
## Draws were sampled using sample(hmc). For each parameter, Bulk_ESS
## and Tail_ESS are effective sample size measures, and Rhat is the potential
## scale reduction factor on split chains (at convergence, Rhat = 1).
```

```
bayestestR::describe_posterior(fit1) #simple summary
```

```
conditional_effects(fit1, resp = "Toscaledlatencytailquiver") #model prediction plots
```

```
conditional_effects(fit1, resp = "Toscaledtailquivercount")
```

```
conditional_effects(fit1, resp = "Toscaledbitepersecondaftersubmitcount")
```

```
conditional_effects(fit1, resp = "Intscaledlatencybite")
```

```
conditional_effects(fit1, resp = "Intscaledbitecount")
```

```
conditional_effects(fit1, resp = "Intproprestagg")
```

```
#checks the data with 10 posterior plots - should overlap nicely
pp_check(fit1, resp = "Toscaledlatencytailquiver")
```

```
pp_check(fit1, resp = "Toscaledtailquivercount")
```

```
pp_check(fit1, resp = "Toscaledbitepersecondaftersubmitcount")
```

```
pp_check(fit1, resp = "Intscaledlatencybite")
```

```
pp_check(fit1, resp = "Intscaledbitecount")
```

```
pp_check(fit1, resp = "Intproprestagg")
```

```
bayes_R2(fit1) #R^2 value of the model
```

```
##                                          Estimate  Est.Error        Q2.5
## R2Toscaledlatencytailquiver             0.2627185 0.10002166 0.066087525
## R2Toscaledtailquivercount               0.1495223 0.08226676 0.017708213
## R2Toscaledbitepersecondaftersubmitcount 0.1025428 0.06626832 0.009865764
## R2Intscaledlatencybite                  0.1718153 0.08551564 0.025084172
## R2Intscaledbitecount                    0.2114177 0.09207547 0.042241864
## R2Intproprestagg                        0.1129019 0.07013392 0.010775912
##                                             Q97.5
## R2Toscaledlatencytailquiver             0.4444277
## R2Toscaledtailquivercount               0.3223895
## R2Toscaledbitepersecondaftersubmitcount 0.2573324
## R2Intscaledlatencybite                  0.3418365
## R2Intscaledbitecount                    0.3876163
## R2Intproprestagg                        0.2718827
```

```
plot(fit1)
```

```
#here, you need to see hairy caterpillars and no trends in the 4 chain outputs

#Plotting the outputs of brms 
#https://cran.r-project.org/web/packages/tidybayes/vignettes/tidy-brms.html#other-visualizations-of-distributions-stat_slabinterval

#7) Plot the model outputs
plot(fit1$fit, pars = c("rescor__Toscaledlatencytailquiver__Toscaledtailquivercount",
                        "rescor__Toscaledlatencytailquiver__Toscaledbitepersecondaftersubmitcount",
                        "rescor__Toscaledtailquivercount__Toscaledbitepersecondaftersubmitcount",
                        "rescor__Intscaledlatencybite__Intscaledbitecount",
                        "rescor__Intscaledlatencybite__Intproprestagg",
                        "rescor__Intscaledbitecount__Intproprestagg",
                        "rescor__Toscaledlatencytailquiver__Intscaledlatencybite",
                        "rescor__Toscaledlatencytailquiver__Intscaledbitecount",
                        "rescor__Toscaledlatencytailquiver__Intproprestagg",
                        "rescor__Toscaledtailquivercount__Intscaledlatencybite",
                        "rescor__Toscaledtailquivercount__Intscaledbitecount",
                        "rescor__Toscaledtailquivercount__Intproprestagg",
                        "rescor__Toscaledbitepersecondaftersubmitcount__Intscaledlatencybite",
                        "rescor__Toscaledbitepersecondaftersubmitcount__Intscaledbitecount",
                        "rescor__Toscaledbitepersecondaftersubmitcount__Intproprestagg"))
```

##### Supplementary information 9: Plotting overt aggression of the same individual across TO and INT

```
# Spearmann correlation: Total aggression across the two roles as TO and INT, without taking into account the response of the other fish.

rho_test8 <- cor.test(focal_to_int$Int_bite_count, focal_to_int$To_bite_count, method = "spearman")

cor8 <- ggplot(focal_to_int, aes(x =To_bite_count, y = Int_bite_count)) +
  geom_point() +  # Add points
  labs(x = "To: Total overt aggression count ", y = "Int: Total overt aggression count")+
  theme_bw() 

cor8 <- cor8 + annotate("text", x = Inf, y = Inf, 
             label = paste("Correlation rho:", round(rho_test8$estimate, 2), 
                           "\n", "p-value:", round(rho_test8$p.value, 4)), 
             hjust = 1.2, vjust = 1.5, size = 4)
cor8
```

```
### Multivariate model: focussing only on the overt aggression in the whole test as TO and INT and how they correlate - not related to competence measures but to see if generally more aggressive individual as a whole is more aggressive as TO and INT in terms of overt aggression - this is a significant correlation in the raw correlations

#1) Get scaled bite counts of Int and To roles
focal_to_int$Int_scaled_bite_count <- as.data.frame(scale(log(focal_to_int$Int_bite_count+1)))[[1]]
focal_to_int$To_scaled_bite_count <- as.data.frame(scale(log(focal_to_int$To_bite_count+1)))[[1]]

#2) Int bite count
bf_Int_bite_count2 <- bf(Int_scaled_bite_count ~ Int_To_scaled_bite_count) + gaussian()

#3) To bite count
bf_To_bite_count2 <- bf(To_scaled_bite_count ~ TO_scaled_int_bite) + gaussian()

#4) Final model is the combination of the two models
bf_total2 <- bf_Int_bite_count2 + bf_To_bite_count2 + set_rescor(rescor = TRUE)

#5) Check the default priors and set good priors, especially for slopes
get_prior(formula = bf_total2, data = focal_to_int)
```

```
#6) Fit and run the brms model
fit2 <- brm(bf_total2,
            data = focal_to_int,
            warmup=1000,
            iter=5000,
            chains=4,
            cores=2,
            backend = "cmdstanr",
            control = list(adapt_delta = 0.90,
                           max_treedepth = 10),
            file="mv_mode_2_")

#7) Adding information criterion to models to use for selection

fit2_1 <- add_criterion(fit2, "loo")

#8) Summary
summary(fit2_1)
```

```
##  Family: MV(gaussian, gaussian) 
##   Links: mu = identity
##          mu = identity 
## Formula: Int_scaled_bite_count ~ Int_To_scaled_bite_count 
##          To_scaled_bite_count ~ TO_scaled_int_bite 
##    Data: focal_to_int (Number of observations: 37) 
##   Draws: 4 chains, each with iter = 5000; warmup = 1000; thin = 1;
##          total post-warmup draws = 16000
## 
## Regression Coefficients:
##                                             Estimate Est.Error l-95% CI
## Intscaledbitecount_Intercept                    0.08      0.17    -0.24
## Toscaledbitecount_Intercept                     0.07      0.17    -0.27
## Intscaledbitecount_Int_To_scaled_bite_count     0.05      0.17    -0.29
## Toscaledbitecount_TO_scaled_int_bite           -0.09      0.17    -0.42
##                                             u-95% CI Rhat Bulk_ESS Tail_ESS
## Intscaledbitecount_Intercept                    0.41 1.00    14279    11024
## Toscaledbitecount_Intercept                     0.42 1.00    12983    11329
## Intscaledbitecount_Int_To_scaled_bite_count     0.39 1.00    13877    10817
## Toscaledbitecount_TO_scaled_int_bite            0.24 1.00    13763    11223
## 
## Further Distributional Parameters:
##                          Estimate Est.Error l-95% CI u-95% CI Rhat Bulk_ESS
## sigma_Intscaledbitecount     1.00      0.13     0.79     1.30 1.00    14832
## sigma_Toscaledbitecount      1.05      0.13     0.83     1.35 1.00    15358
##                          Tail_ESS
## sigma_Intscaledbitecount    11033
## sigma_Toscaledbitecount     12400
## 
## Residual Correlations: 
##                                              Estimate Est.Error l-95% CI
## rescor(Intscaledbitecount,Toscaledbitecount)     0.34      0.16    -0.01
##                                              u-95% CI Rhat Bulk_ESS Tail_ESS
## rescor(Intscaledbitecount,Toscaledbitecount)     0.62 1.00    11210    11233
## 
## Draws were sampled using sample(hmc). For each parameter, Bulk_ESS
## and Tail_ESS are effective sample size measures, and Rhat is the potential
## scale reduction factor on split chains (at convergence, Rhat = 1).
```

```
bayestestR::describe_posterior(fit2) #simple summary
```

```
# check fit - hairy caterpillar plot
plot(fit2)
```

```
# ppcheck
pp_check(fit2, resp = "Intscaledbitecount")
```

```
pp_check(fit2, resp = "Toscaledbitecount")
```

```
# Plot the residual correlations
plot(fit2$fit, pars = c("rescor__Intscaledbitecount__Toscaledbitecount"))
```

```
rescor <- as_draws_df(fit2)

rescor_value <- rescor$rescor__Intscaledbitecount__Toscaledbitecount

df <- data.frame(correlation = rescor_value)

library(ggplot2)

rescor_value <- data.frame(
  estimate = 0.34,
  lower = -0.01,
  upper = 0.62
)

ggplot(df, aes(x = correlation)) +
  geom_density() +
  geom_vline(
    xintercept = rescor_value$estimate,
    linetype = "dashed"
  ) +
  annotate(
    "text",
    x = Inf,
    y = Inf,
    label = paste0(
      "Estimate = ", rescor_value$estimate,
      "\n95% CI = [", rescor_value$lower,
      ", ", rescor_value$upper, "]"
    ),
    hjust = 1.1,
    vjust = 1.5
  ) +
  theme_classic()
```
